# A Degron Decoy System Co-opts Pathological Seeding to Enable Clearance of Multimeric α-Synuclein

**DOI:** 10.64898/2026.02.23.706428

**Authors:** Gillian E. Gadbois, Alexander P. Plonski, Galia T. Debolouchina, Fleur M. Ferguson

## Abstract

Pathological seeding of protein misfolding is a hallmark of proteinopathies. However therapeutic strategies to clear these aggregates are lacking, impairing both study of their biological importance in disease etiology and progression as well as development of therapeutics. This is due in part to the need to selectively clear oligomerized proteins whilst leaving functional monomers intact, as well as the challenge of developing molecules that act on the full complement of ‘misfolds’ the protein can adopt throughout the course of disease. In this work, we describe a dopant system consisting of an engineered alpha-synuclein protein construct that rapidly co-aggregates into existing WT alpha-synuclein oligomers, enabling rapid degradation of the entire assembly in the presence of a small molecule trigger. This work provides proof-of-principle for an approach that transforms pathological seeding from a disease-driver into a therapeutic vulnerability, and is potentially applicable to any proteinopathy without requiring a small molecule binder of the pathologic species.

## INTRODUCTION

The misfolding and accumulation of alpha-synuclein in the brain is a hallmark of Lewy Body Dementias, that include Parkinson’s Disease and Dementia with Lewy Bodies.^1,2^ These disorders affect approximately 1.3 million people in the US, and are characterized by decline in the autonomic and central nervous systems that lead to defects in motor function and cognition, and to hallucinations and REM-sleep disorders.^3,4^ No disease-modifying therapies are available for Lewy Body Dementia’s, with the standard of care levodopa working as a palliative therapy to boost levels of dopamine and thus relieve symptoms in patients.^5–7^

To better understand the etiology of Lewy Body Dementias, the conformations adopted by misfolded alpha-synuclein species have been rigorously characterized by the neuroscience and structural biology communities.^8^ Here, remarkable polymorphism has been observed, with different aggregate structures detected dependent on patient^9^ and disease status,^10^ pH^11^ and salt concentration^12^, structure of the “seed” used to nucleate the aggregate^13^, and the post-translational modification state of alpha-synuclein.^14^ These different ‘strains’ are reported to have different biological properties such as toxicity and seeding capacity.^15^ There are likely other yet-to-be discovered factors that influence alpha-synuclein aggregate structure. Unsurprisingly, given the structural heterogeneity of misfolded alpha-synuclein species, efforts to develop positron-emission tomography (PET) tracers that selectively bind to pathogenic alpha-synuclein proteoforms have yet to yield molecules suitable for clinical diagnostic application.^16,17^ Instead, the FDA supports the use of seed amplification assays, that measure the capacity of alpha-synuclein species from a patient to trigger aggregation of recombinant alpha-synuclein, in both clinical and research studies.^18,19^ This differentiates alpha-synuclein from other proteinopathies such as tauopathies, as the diagnostic criteria measures the seeding function, rather than the fold (PET tracer binding), to define disease-associated proteoforms.^20–22^

The role of post-translational alpha-synuclein phosphorylation at serine 129 has also been investigated for its role in pathogenesis. Alpha-synuclein pS129 is found in over 80% of patient brains with Lewy Body Dementia, and less than 5% of healthy brains, a clear association with disease state.^23^ Recent work suggests that phosphorylation at this site is a protective mechanism, that occurs following misfolding and soluble oligomer formation, to slow the aggregation formation process.^24,25^ Hence, in cellular models of Parkinsons’ Disease, pS129 alpha synuclein is used as a marker of oligomeric species in the soluble cellular fraction. These soluble alpha-synuclein oligomers represent intermediates between healthy and insoluble aggregate protein conformations.^26^ Emerging evidence points to oligomers as the primary pathogenic species that cause the lion’s share of membrane damage, seeding, propagation, and toxicity in Parkinsons Disease.^27^

Clearance of misfolded or aggregated alpha-synuclein in patients is one of several therapeutic hypotheses being pursued in efforts to slow or halt disease progression. To this end, bifunctional targeted protein degraders that can clear aggregated alpha synuclein have been developed.^28–32^ These small molecules use pan-aggregate binding pharmacophores, such as benzothiazoles, as the misfolded alpha-synuclein recruiter, linked to binders of E3-ligases or autophagy-targeting chaperones via a chemical spacer. Reported alpha-synuclein degraders have shown degradation activity in model systems such as HEK293 cells or rat neurons overexpressing human alpha-synuclein, where aggregation is seeded by addition of recombinant alpha-synuclein preformed fibrils (PFFs).^28–32^ These small molecule degraders represent an exciting advance in our quest to better understand and potentially treat Lewy Body Dementias.

However, existing alpha-synuclein degraders have limitations that have hindered their wide use in research. The primary limitation of current small molecule alpha-synuclein degraders is a dependence on the model system expressing a ligandable alpha-synuclein aggregate species that binds the warhead incorporated in the small molecule degrader. As detailed above, there is a paucity of potent, selective, clinically-validated binders of misfolded alpha-synuclein, and unlike other proteinopathy aggregates that have been targeted by bifunctional small molecule degraders, such as misfolded tau,^33,34^ tracer binding is not part of the diagnostic criteria for synucleinopathies, which are instead defined by their ability to seed co-aggregation of monomeric alpha-synuclein.^21^ Further, the misfolded protein species present in a model system of interest may vary, due to batch-to-batch variability in the PFFs, variability in the seeding conditions, and many other factors listed above, leading to reduced robustness of these degraders activity across different model systems.^35^ Crucially, soluble oligomeric pS129 alpha-synuclein species are not universally targetable by published alpha-synuclein degrader molecules. This is particularly important as in most reported patient iPSC-derived dopaminergic neuron models of synucleinopathies, ThT positive aggregates are not detectable even at time points as late as 90 days post differentiation, but soluble pS129 oligomeric species are present.^36^ Alpha-synuclein also represents an exceptionally high-bar for targeted protein degradation, as it is one of the most highly expressed proteins in healthy brain, representing ∼ 1% of the total proteome, and these protein levels are further elevated in Parkinsons’ Disease.^37–40^ Many currently disclosed small molecule bifunctional degraders of alpha-synuclein have limitations, including slow degradation kinetics, weak potency, and unknown global selectivity for their targets, highlighting a need for development of orthogonal approaches.

These limitations motivated us to create a complementary set of tools with which to modulate pathogenic alpha-synuclein species in living cells. To do so, we sought to leverage the defining pathogenic features of LBD-associated alpha-synuclein, namely its propensity to oligomerize soluble alpha-synuclein monomers. Here, we report a chemical-genetic degradation system, that leverages the seeding activity of misfolded alpha-synuclein to enable incorporation of a carefully designed degron-tagged alpha-synuclein decoy construct in living cells and neurons. Addition of a small molecule degrader enables recruitment of the E3-ligase CRL4^CRBN^ to the multimeric assembly, triggering ubiquitination that leads to rapid, selective, degradation of all aggregates and soluble pS129 positive alpha-synuclein oligomers via a bridged or collateral degradation mechanism. Crucially, soluble, non-pathogenic, endogenous alpha-synuclein proteins remain unaffected, whilst excess tagged construct is rapidly cleared, creating a traceless system.

## RESULTS

To investigate whether we could degrade seeding-competent alpha-synuclein via a degron-tagged dopant we designed alpha-synuclein degron tag fusions. To develop a degron-tagged alpha-synuclein construct compatible with our approach, we sought both rapid co-aggregation kinetics and rapid degradation kinetics. We designed alpha-synuclein degron tag fusions that incorporated alpha-synuclein with an A53T mutation, a disease associated mutation that has shown to increase misfolding of the protein. For tags, we turned to the immunomodulatory drug (IMiD) inducible tags that enable proteasomal degradation of the tagged protein via the recruitment of the E3-ligase substrate adapter protein cereblon (Figure 1A).^41,42^ These tags are small in size, the Super degron is 60 amino acids long^41^ and the Minimal degron is 23 amino acids long^43^, an important consideration as alpha-synuclein is only 160 amino acids (15 KDa), and inclusion of a larger tag may impair the ability of the fusion construct to be seeded by and co-aggregate with wild-type alpha-synuclein species, or may be too far from the oligomeric core to trigger collateral degradation of wild-type co-aggregated alpha-synuclein. In addition, these tag-systems have been optimized to work with IMiDs designed to have minimal off-targets.^44^ Unlike many other degron tag systems based on bifunctional molecules, the IMiD drugs and analogs have favorable *in vivo* pharmacokinetics and are blood-brain barrier permeable, an important consideration when studying CNS-targets.^45,46^ To determine the optimal tag composition and location, we designed Super degron N-terminal (Super-SNCA^A53T^), Super degron C-terminal (SNCA^A53T^-Super), Minimal degron N-terminal (Min-SNCA^A53T^), and Minimal degron C-terminal constructs (SNCA^A53T^-Min). All the constructs contain an HA tag to facilitate differentiation of the degron tagged protein from the wild-type protein in co-expression systems and a (GGGGS)_2_ spacer (Figure S1). To induce degradation, we used the IMiD, pomalidomide (POM), and a second-generation analogue, compound 27 (BRD1155), designed to degrade fewer endogenous zinc finger targets (Figure 1B).^44^

**Figure 1.**
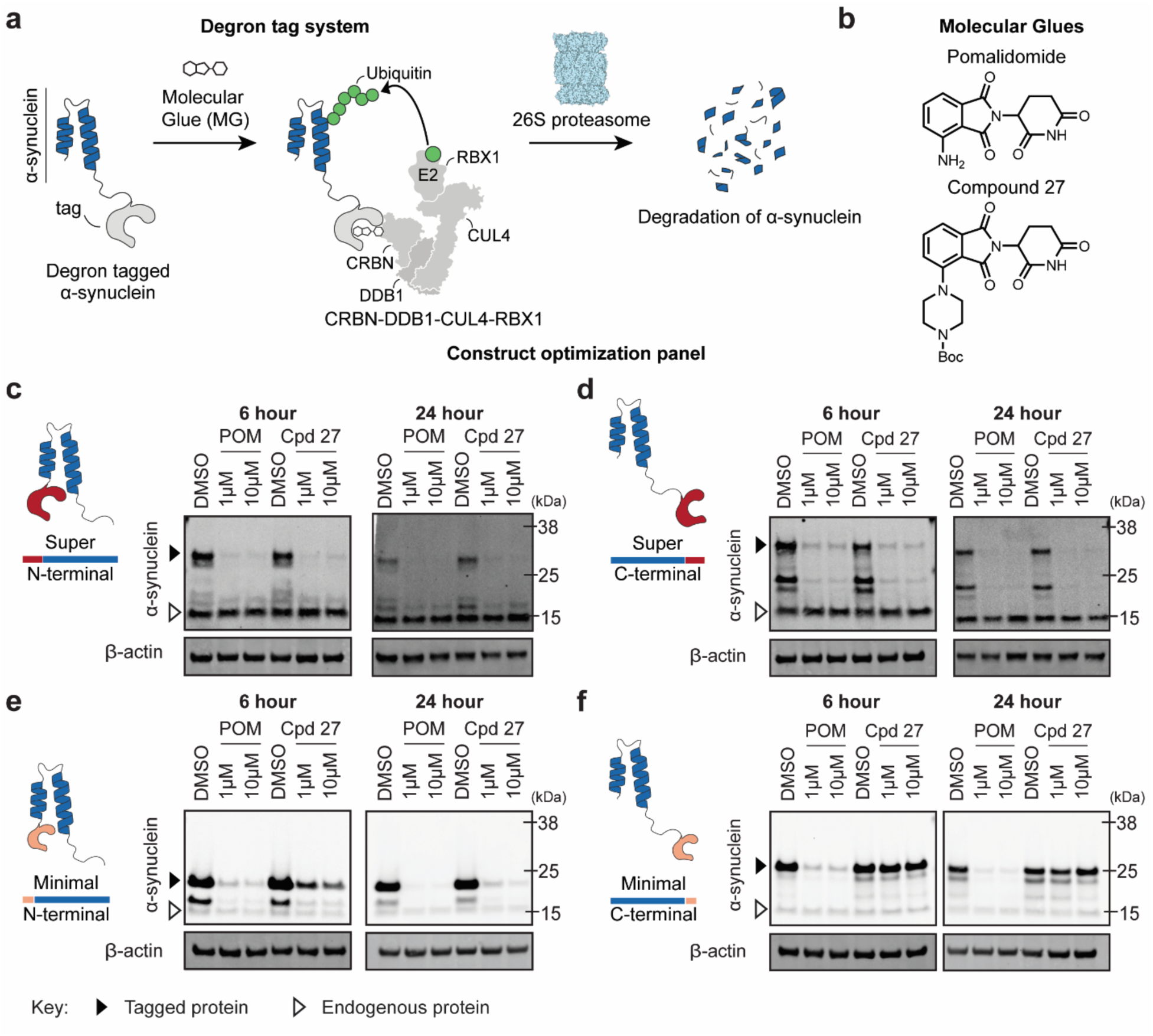
Degron tags enable degradation of soluble alpha-synuclein via small molecules. **a.** Schematic representation of degradation mechanism. **b.** Chemical structures of pomalidomide (POM) and compound 27 (BRD1155). **c-f.** Western blots of alpha-synuclein levels following 6- or 24-hour treatment of pomalidomide or compound 27 in cells expressing degron tagged A53T alpha-synuclein. **c.** Super degron N-terminal tag (Super-SNCA^A53T^). **d.** Super degron C-terminal tag (SNCA^A53T^-Super) **e.** Minimal degron N-terminal tag (Min-SNCA^A53T^). **f.** Minimal degron C-terminal tag (SNCA^A53T^-Min). Data in c-f representative of *n* = 3 independent experiments.

We first set out to evaluate if the soluble form of each tagged construct could be rapidly and completely degraded by addition of Pomalidomide or Cpd 27. To determine the ability of the degron constructs to degrade alpha-synuclein, we performed western blot analysis of alpha-synuclein in HEK293 cell lines stably overexpressing the degron constructs, treated with Pomalidomide or Cpd 27 for 6 hours or 24 hours. The Super-SNCA^A53T^ and SNCA^A53T^-Super constructs showed significant degradation with both compounds at 6 hours and complete degradation at 24 hours (Figure 1C,D). Min-SNCA^A53T^ enabled significant degradation with Pomalidomide at 6 hours and complete degradation at 24 hours, while Cpd 27 showed minimal degradation at 6 hours and enhanced degradation at 24 hours (Figure 1E). SNCA^A53T^-Min showed significant degradation at 6 hours with Pomalidomide and complete degradation at 24 hours but no degradation with Cpd 27 at 6 or 24 hour time points (Figure 1F). In all conditions, wild-type (non-tagged) alpha-synuclein levels were not affected. We selected Min-SNCA^A53T^ and SNCA^A53T^-Min for further evaluation based on their smaller tag size.

To evaluate the selectivity of the degradation, we performed global proteomics analysis in cells stably expressing Min-SNCA^A53T^ or SNCA^A53T^-Min constructs treated with 1 µM of Pomalidomide or Cpd 27 for 6 hours (Figure S2). No off-target degradation was observed in the cells expressing Min-SNCA^A53T^(Figure S2 A,B), while modest degradation of RAB28 and ZFP91 was observed in the SNCA^A53T^-Min expressing cells (Figure S2C). We observed significant degradation of alpha-synuclein following treatment with either Pomalidomide or Cpd 27 in cells expressing Min-SNCA^A53T^ (Figure S2A,B), but only observed significant degradation of alpha-synuclein following Pomalidomide treatment, but not Cpd 27 treatment, in cells expressing SNCA^A53T^-Min (Figure S2C,D). These findings align well with the western blot results and further support that the degron tags can enable rapid, selective proteasomal degradation of tagged alpha-synuclein. This potent and selective degradation of the soluble monomeric versions of the tagged constructs is important, as it allows for rapid removal of construct that does not co-aggregate without altering wild-type healthy alpha-synuclein levels (Figure 1E). Based on these data, we selected the Min-SNCA^A53T^ construct due to its small size, robust expression, and rapid, selective and complete degradation by both Pomalidomide and Cpd 27.

To evaluate if the degron tagged alpha-synuclein could co-aggregate with endogenous misfolded alpha-synuclein, we recombinantly expressed both WT SNCA and Min-SNCA^A53T^ proteins (Figure S3). The proteins’ identity was confirmed by intact mass TOFMS using an Agilent 1260 Infinity Binary LC coupled to a 6230 TOFMS system (Figure S4-5). Following purification, we aggregated each protein using a 2-week agitation protocol, as either a pure sample, or a mixed Min-SNCA^A53T^:WT sample.

We measured the *in vitro* aggregation kinetics of Min-SNCA^A53T^ alone, or when seeded by WT alpha-synuclein pre-formed fibrils using a ThT fluorescent reporter assay (Figure 2A). We observed the desired rapid and dose-dependent templated co-aggregation upon addition of WT alpha-synuclein pre-formed fibrils (PFFs), and slow aggregation in the absence of a seeding species (Figure 2B,C). To verify the morphology of the Min-SNCA^A53T^ aggregates and co-aggregates were consistent with those formed by pure WT protein, we performed transmission electron microscopy (TEM) (Figure 2D, Figure S6). Here, we observed broadly uniform aggregate morphology across the samples, indicating that aggregate morphology is faithfully preserved in mixed population aggregates.

**Figure 2.**
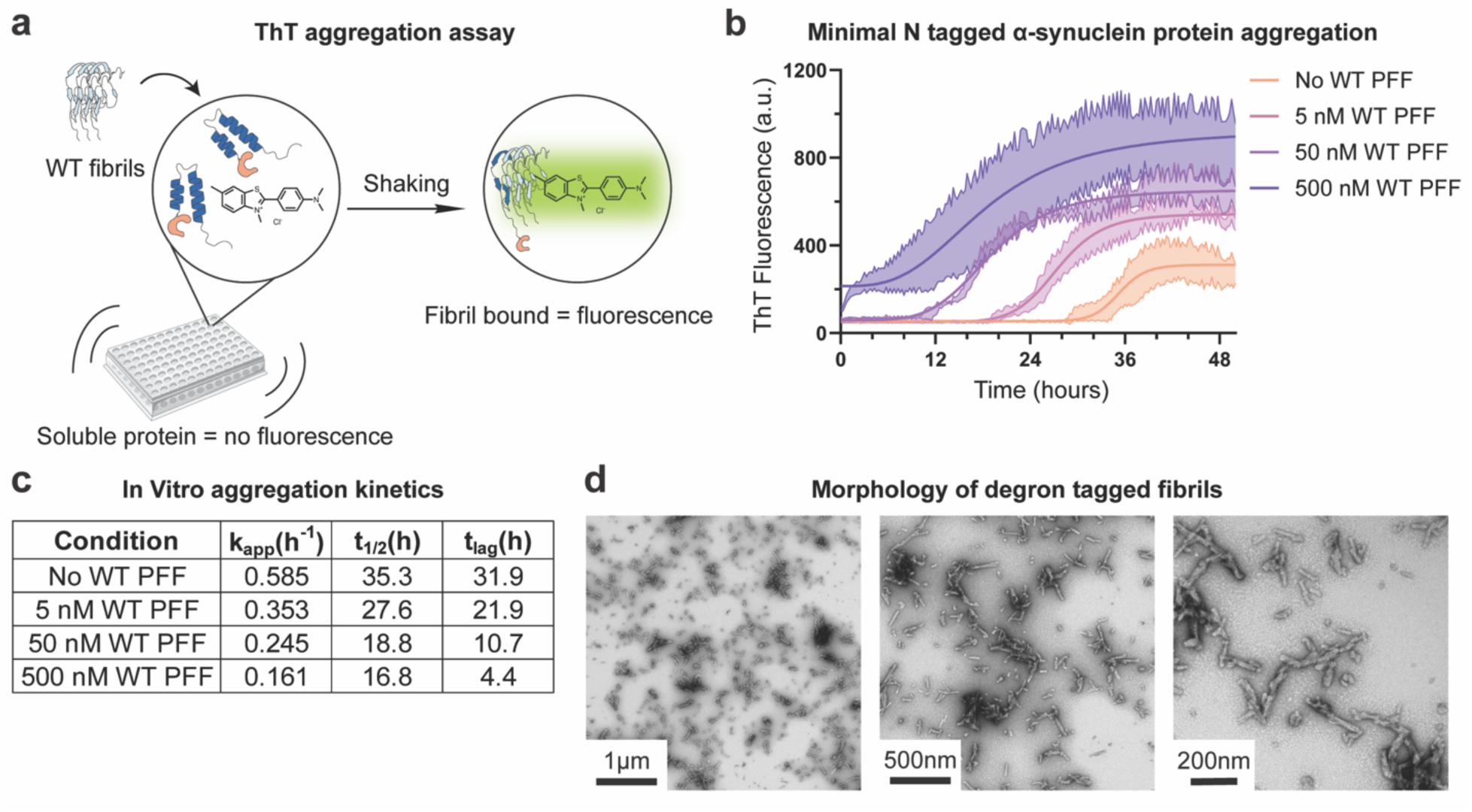
Minimal degron tagged alpha-synuclein can aggregate into fibril. **a.** Scheme of in vitro ThT aggregation assay. **b.** ThT fluorescence of Min-SNCA^A53T^ protein seeded with untagged wild type alpha-synuclein preformed fibrils (PFFs). **c.** Kinetic parameters from ThT aggregation assay. **d.** Transmission electron microscopy of N-terminal minimal degron tagged alpha-synuclein fibrils. Data in b presented as average +/- s.d. of *n* = 3 technical replicates.

To compare the ability of our degron-tagged construct to co-aggregate with WT alpha-synuclein fibrils in living cells, we stably expressed all candidate degron constructs, alongside endogenous alpha-synuclein in HEK293 cells, and then treated them with sonicated WT PFFs to seed co-aggregation (Figure 3A). Following 24 hours of co-aggregation, cells were treated with DMSO, or with 1-10 µM Pomalidomide or Cpd 27 and degradation was evaluated by both immunoblot and global proteomics analysis following extraction in 3% SDS containing lysis buffer. We used denaturing gel electrophoresis to separate solubilized alpha-synuclein species from detergent-resistant multimers.

**Figure 3.**
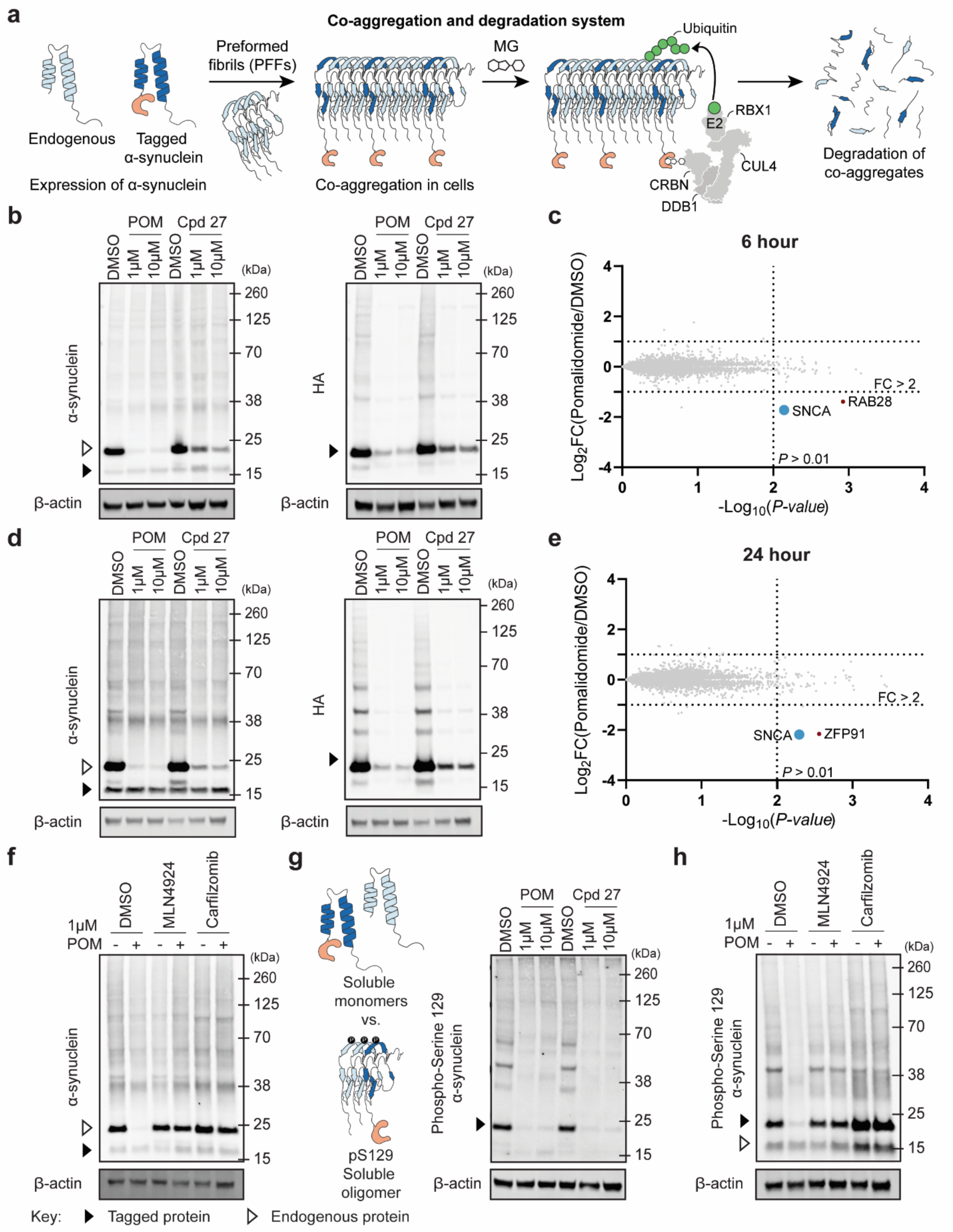
Clerance of preformed fibril aggregated alpha-synuclein using minimal degron tags. **a.** Schematic representation of preformed fibril induced co-aggregation and degradation of alpha-synuclein. **b.** Western blots of alpha-synuclein and HA levels following 6-hour treatment of pomalidomide or compound 27 in Min-SNCA^A53T^ expressing cells transfected with PFFs. HA tag on degron-tagged alpha-synuclein. **c.** Global proteomics of Min-SNCA^A53T^ expressing cells transfected with PFFs and treated with 1 µM pomalidomide for 6 hours. **d.** Western blots of alpha-synuclein and HA levels following 24-hour treatment of pomalidomide or compound 27 (BRD1155) in Min-SNCA^A53T^ expressing cells transfected with PFFs. HA tag on degron-tagged alpha-synuclein. **e.** Global proteomics of Min-SNCA^A53T^ expressing cells treated with PFFs followed by 1 µM pomalidomide for 24 hours. **f.** Degradation is rescued by pre-treatment of 1 µM MLN4924 or 0.25 µM Carfilzomib. **g**. Degradation results in clearance of phospho-Serine 129 alpha-synuclein, a marker of disease relevant oligomers. **h.** Degradation of disease relevant oligomers is rescued by pre-treatment of 1 µM MLN4924 or 0.25 µM Carfilzomib. **f-h**. Min-SNCA^A53T^ expressing cells treated with PFFs followed by 1 µM pomalidomide for 24 hours FC = fold change. Data in b, d, f-h representative of *n* = 3 independent experiments. Data in c,e calculated from *n* = 3 technical replicates.

At the 6-hour time point, we observed modest levels of co-aggregation via the presence of higher molecular weight bands and subsequent degradation in the Min-SNCA^A53T^ expressing cell line following compound treatment (Figure 3B). By the 24-hour time point, we observed clear co-aggregation and potent degradation in the Min-SNCA^A53T^ expressing cell line with both Pomalidomide and Cpd 27 (Figure 3D). We determined the proteome-wide selectivity of the co-aggregate degradation induced by Pomalidomide and Cpd 27 in WT alpha-synuclein PFF-treated cells expressing Min-SNCA^A53T^ alongside endogenous alpha-synuclein using mass-spectrometry based global proteomics analysis. Here, we observed degradation of alpha-synuclein with Pomalidomide after both 6 hours and 24 hours treatment (Figure 3C,E). RAB28 and ZFP91 were again degraded as off-targets in a subset of conditions. These findings indicate that the Minimal N-terminal tag system enables co-aggregation and rapid, selective degradation of alpha-synuclein multimers.

In contrast to the Min-SNCA^A53T^ system, we observed some co-aggregation with the SNCA^A53T^-Min expressing line but only observed degradation with Pomalidomide and not Cpd 27 (Figure S7). Neither of the Super degron tagged constructs co-aggregated following treatment with the PFFs, likely due to the larger tag size (Figure S8).

To demonstrate that the degradation of alpha-synuclein proceeds with the proposed mechanism, we performed rescue experiments, pretreating Min-SNCA^A53T^ expressing cells with MLN4924, an inhibitor of NAE-mediated cullin-ring ligase activation, or carfilzomib, a proteasome inhibitor, and compound treatment. We observed complete rescue of total alpha-synuclein levels by immunoblot supporting a targeted protein degradation mechanism (Figure 3F).

To evaluate the ability of our degron-based system to clear the oligomeric alpha-synuclein marked by pS129, we treated cells stably expressing Min-SNCA^A53T^ alongside endogenous alpha-synuclein with sonicated WT alpha-synuclein PFFs for 24 hours to seed co-aggregation. We treated cells with DMSO, Pomalidomide or Cpd 27 for 6 hours or 24 hours, and blotted for pS129 (Figure 3G). Here, we observed complete clearance of both the detergent-soluble pS129 positive alpha-synuclein species, as well as the higher molecular weight detergent-resistant pS129 bands upon treatment with degraders. To demonstrate that this degradation is on-mechanism, we performed rescue experiments, pretreating cells with MLN4924 or carfilzomib, and observed complete rescue of pS129 alpha-synuclein degradation (Figure 3H).

Taken together, these data suggest that tag size and location play a profound role in the ability of alpha-synuclein to aggregate following seeding, and for the degron tag to be accessible for degradation when part of the aggregate. We concluded the Min-SNCA^A53T^ construct as the most optimal for continued validation.

Finally, we sought to validate our Min-SNCA^A53T^ system in a more disease-relevant model of Parkinsons’ Disease. To do so, we transfected SHSY-5Y cells with lentiviral constructs to enable expression of Min-SNCA^A53T^ alongside WT endogenous alpha-synuclein, and differentiated them into neurons following established protocols (Figure 4A). We evaluated three different expression levels; a CMV promoter to promote high levels of degron-tagged construct expression, a synaptin promoter to ensure physiologically relevant lower levels of expression, and a tet-inducible promoter to mimic a therapeutic setting where degron-tagged Min-SNCA^A53T^ construct expression occurs in mature neurons.

**Figure 4.**
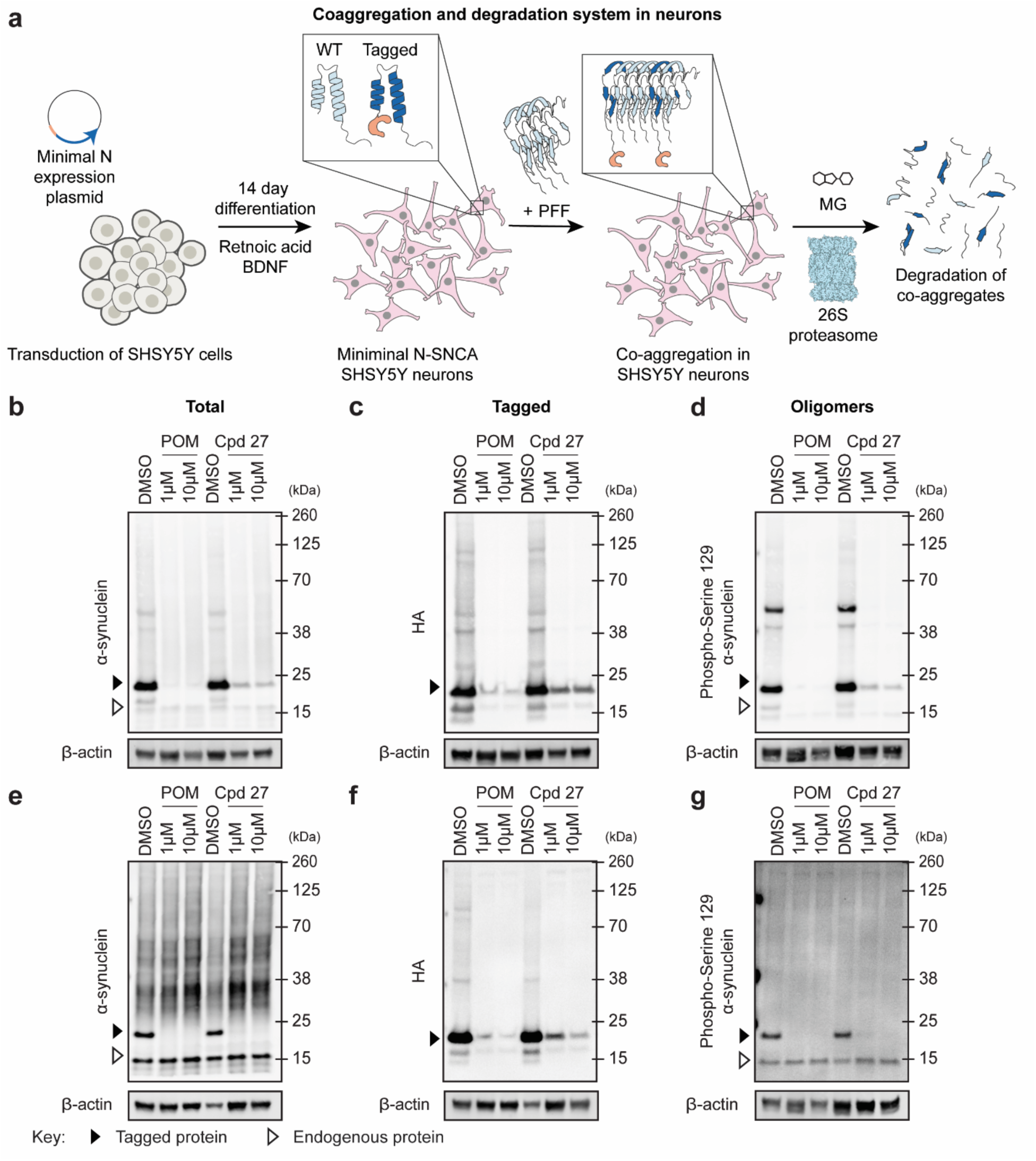
Degron tag system enables co-aggregation and degradation of alpha-synuclein in neurons. **a.** Schematic representation of differentiation and treatment strategy. **b-d.** Western blots of SHSY5Y neurons expressing Min-SNCA^A53T^ via CAG promoter treated with PFFs for 24 hrs, followed by 24-hour compound treatment. **b.** Total alpha-synuclein. **c.** Degron tagged alpha-synuclein. **d.** Disease relevant pS129 oligomers. **e-g.** Western blots of SHSY5Y neurons expressing Min-SNCA^A53T^ via DOX inducible promoter treated with preformed fibrils, followed by 24-hour compound treatment. **e.** Total alpha-synuclein. **f.** Degron tagged alpha-synuclein. **g.** Disease relevant oligomers. Data in b-g representative of *n* = 3 independent experiments.

We treated the neurons with sonicated WT alpha-synuclein PFFs to induce co-aggregation for 24 hours. Next, we treated cells with DMSO, Pomalidomide or Cpd 27. Gratifyingly, we observed efficient co-aggregation of WT and degron-tagged alpha-synuclein expressed via a the synaptin promoter. This led to efficient degradation upon treatment with Pomalidomide or Cpd 27 of soluble degron-tagged alpha-synuclein, mixed oligomers, and mixed aggregates, while untagged soluble alpha-synuclein was unchanged (Figure 4B-D). In contrast, use of a CMV promoter led to reduced co-aggregation and clearance, indicating that high levels of degron-tagged construct are not required for, and may be detrimental to, mixed aggregate clearance (Fig. S9). Next, we induced expression of Min-SNCA^A53T^ in mature neurons for 24 hours using doxycycline, followed by treatment with sonicated WT alpha-synuclein PFF for 24 hours, and then compound treatment for 6 hours or 24 hours. We also observed co-aggregation and efficient clearance of alpha-synuclein species under these conditions (Figure 4E-G).

## DISCUSSION

Here we describe a degron-tagged dopant approach that exploits the pathogenic features of alpha-synuclein, namely its propensity to oligomerize and seed co-aggregation, to enable targeted protein degradation of endogenous alpha-synuclein aggregates and soluble oligomers, while leaving soluble WT alpha-synuclein untouched. This simple but powerful system enables selective targeting of misfolded species, while circumventing the requirement for selective tracer development, a major challenge in the field. In this manuscript, we apply the system to degrade aggregated and oligomeric alpha-synuclein as a proof-of-principle, however, our system could be theoretically applied to any proteinopathy-associated protein.

Our system provides a complementary approach to existing chemical genetic tools that have been reported for degradation of misfolded proteins. For example, TRIMTACs enable degradation of oligomeric proteins via the use of a TRIM21 tag, where protein oligomerization triggers TRIM21 E3-ligase activation, and subsequent proteasomal degradation.^47–49^ In this system, many subunits in the oligomer must contain a tag to effect oligomerization-mediated TRIM21 activation, allowing for selective degradation based on proteoform, but favoring a knock-in or high expression level of the tag. Our system allows for transient or substochiometric inclusion of the tagged protein, widening applications to study of endogenous aggregates and model systems and reducing the time-intensive process of generating knock-in cells and mice. In addition, because these systems are state-specific, the non-oligomerized tagged monomer remains in the system. In our system, soluble tagged protein (but not soluble untagged endogenous protein) is degraded rapidly upon small molecule addition providing a traceless background.

A more recent iteration, whereby a TRIM21 recruiting ligand is used to recruit TRIM21 to oligomers has been described. While a promising approach, it has so far been deployed against tagged tau proteins to prevent templated aggregation.^49^ Our system offers a complementary functionality, by allowing for clearance of *existing* oligomers.

The AdPROM system has also been deployed against misfolded proteins. In this system, a biological degrader construct composed of the VHL E3-ligase substrate adaptor protein linked to a nanobody of alpha-synuclein has been used to globally reduce alpha-synuclein levels in the cell.^50^ Our system complements this system, as it only targets oligomeric or aggregated WT synuclein, and is inducible via addition of a small molecule, adding temporal control.

Limitations of our system include the required careful optimization of the tagged construct to ensure it is compatible with co-aggregation induced by the endogenous, WT misfolded protein seeds and to ensure the resultant co-aggregates can enable the collateral degradation of the entire composite oligomer. A second limitation is the incompatibility with murine model systems, due to species-specific differences in CRBN that render the glue molecules unable to promote degradation in that context. Here, workarounds such as working in model systems that express humanized CRBN, or use of alternative tags that are compatible with murine CRBN^51^, may be possible, but add another layer of engineering to the approach.

Potential future investigation includes the generalization of the platform across unliganded misfolded proteins, such as FUS and TDP-43, as well as translational work to develop mRNA or protein based *in vivo* tools for therapeutic intervention.

## Supporting information

Uncropped Blots

## Acknowledgements

This work was supported by the Larry L. Hillblom Foundation Grant #2022-A-008-SUP (F.M.F.). This work was supported by NIH P01AG081167 (G.T.D.) and DP2NS132610 (F.M.F). G.E.G. and A.P.P. were supported by the NIH Molecular Biophysics Training Grant T32GM139795. Lentivirus was produced by Sanford Burnham Prebys Viral Vector core (Shared Instrumentation Grant S10 OD036254). We would like to thank Chun-Teng Huang for his assistance. Proteomic studies were performed at the UC San Diego Goeddel Family Sandbox. We would like to thank Dr. Fulin Jiang for his assistance. We would like to thank the University of California, San Diego - Cellular and Molecular Medicine Electron Microscopy Core (UCSD-CMM-EM Core, RRID: SCR_022039) for providing access to equipment and technical assistance for the TEM imaging. We would like to thank Amit Choudhary and Sreekanth Vedagopuram for their assistance with construct design and providing compound 27 (BRD1155), funded by NIH R01GM137606.

---

**Supplementary Figure 1.**
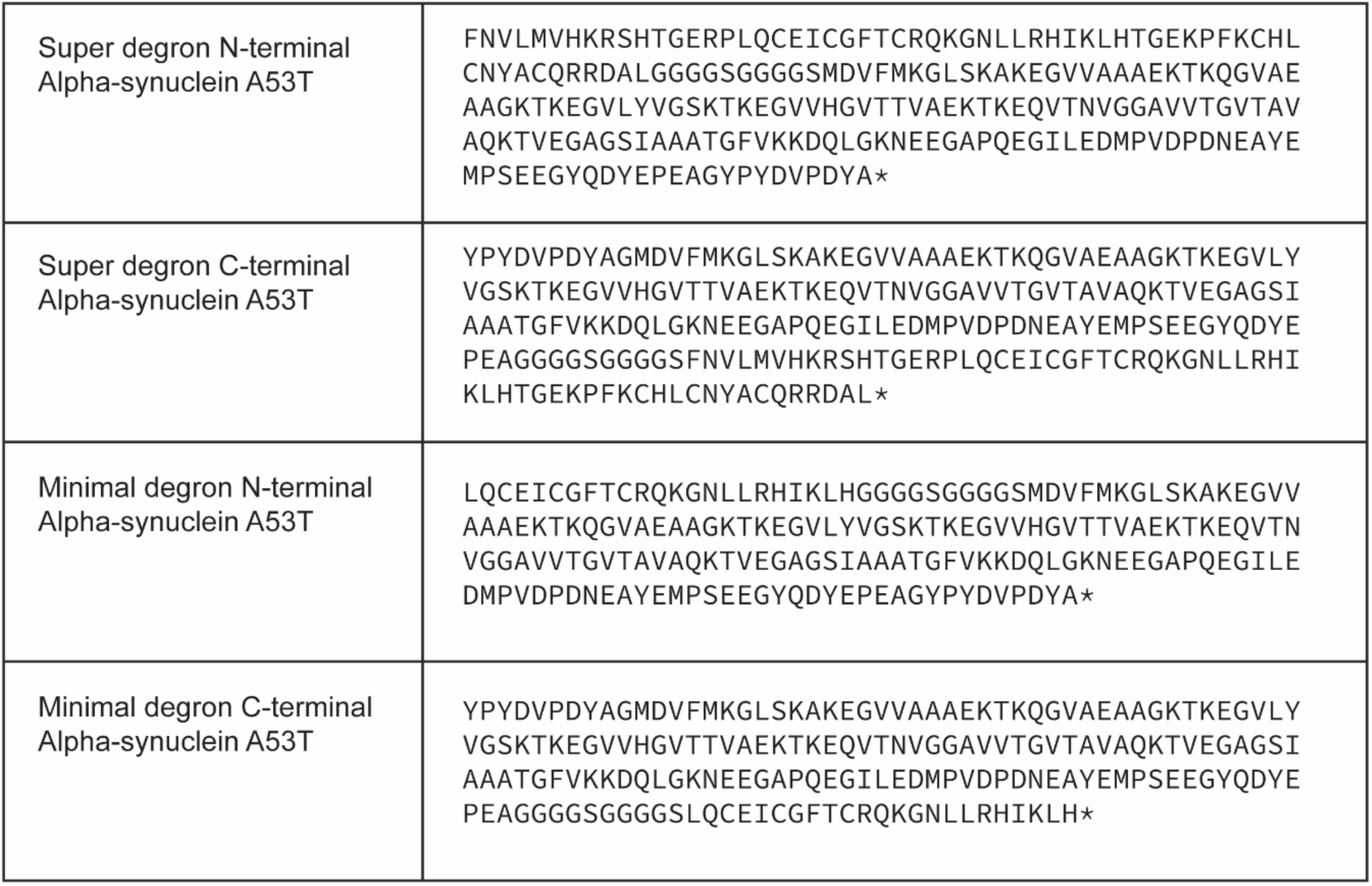
Sequence of degron tagged A53T alpha-synuclein constructs including HA tag and linkers.

**Supplementary Figure 2.**
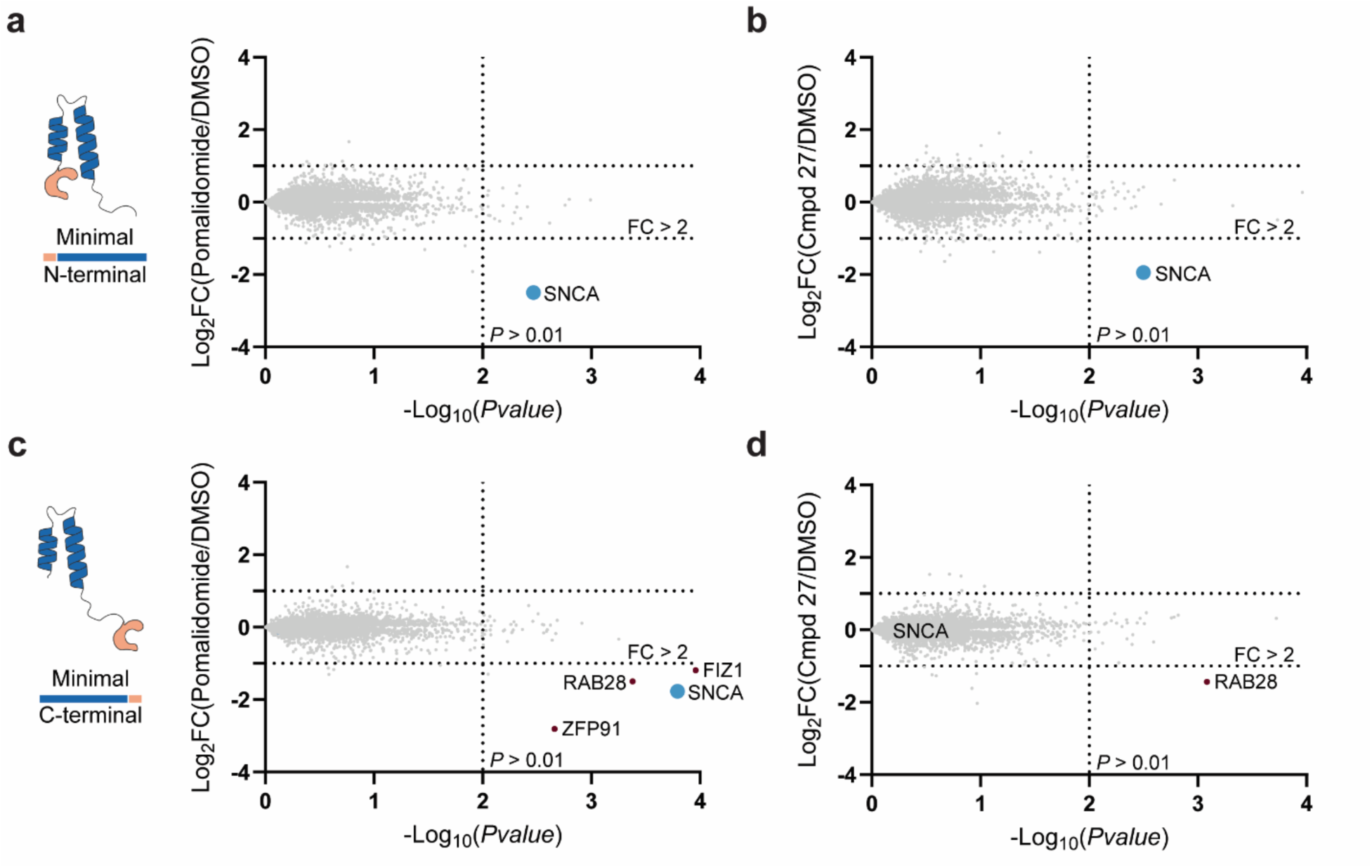
Selectivity profile of degron tag enabled degradation of alpha-synuclein. **a-b.** Global proteomics of Min-SNCA^A53T^ expressing cells treated with **a.** 1 µM pomalidomide or **b.** 1 µM compound 27 for 6 hours. **c-d.** Global proteomics of SNCA^A53T^-Min expressing cells treated with **c.** 1 µM pomalidomide or **d.** 1 µM compound 27 for 6 hours. FC = fold change.

**Supplementary Figure 3.**
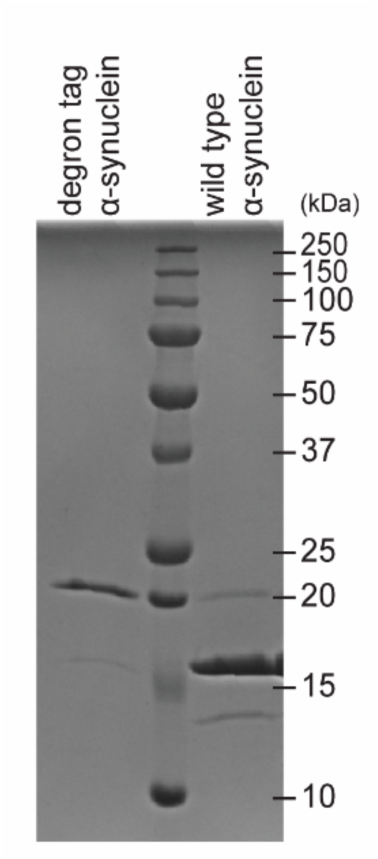
Protein post-HPLC purification.

**Supplementary Figure 4.**
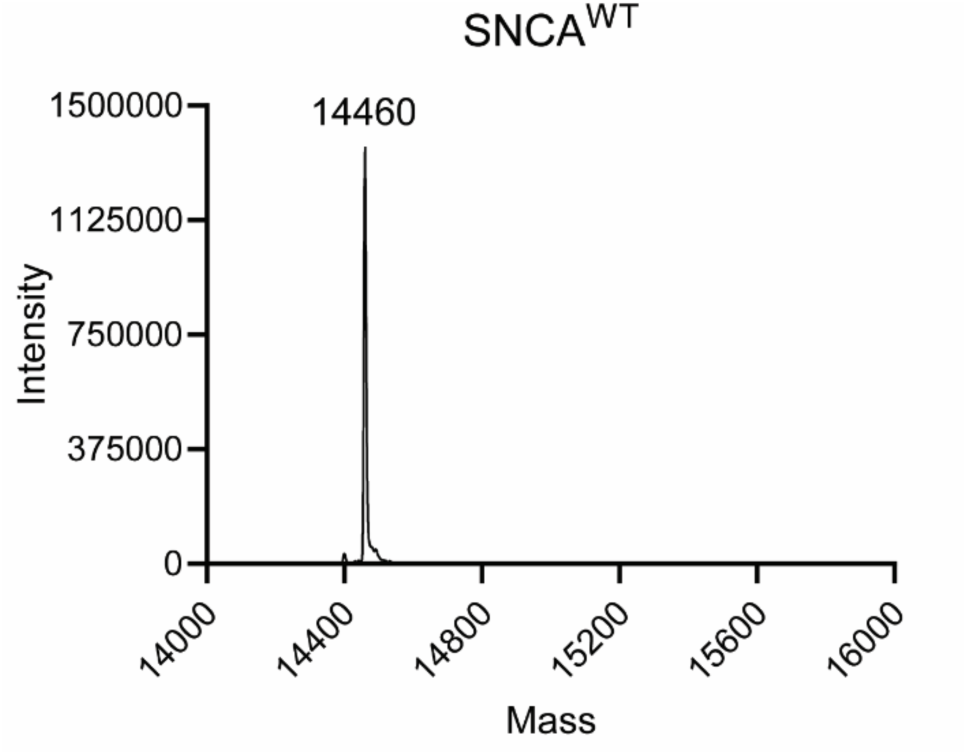
Mass spectrometry analysis of purified wild-type alpha-synuclein (SNCA^WT^).

**Supplementary Figure 5.**
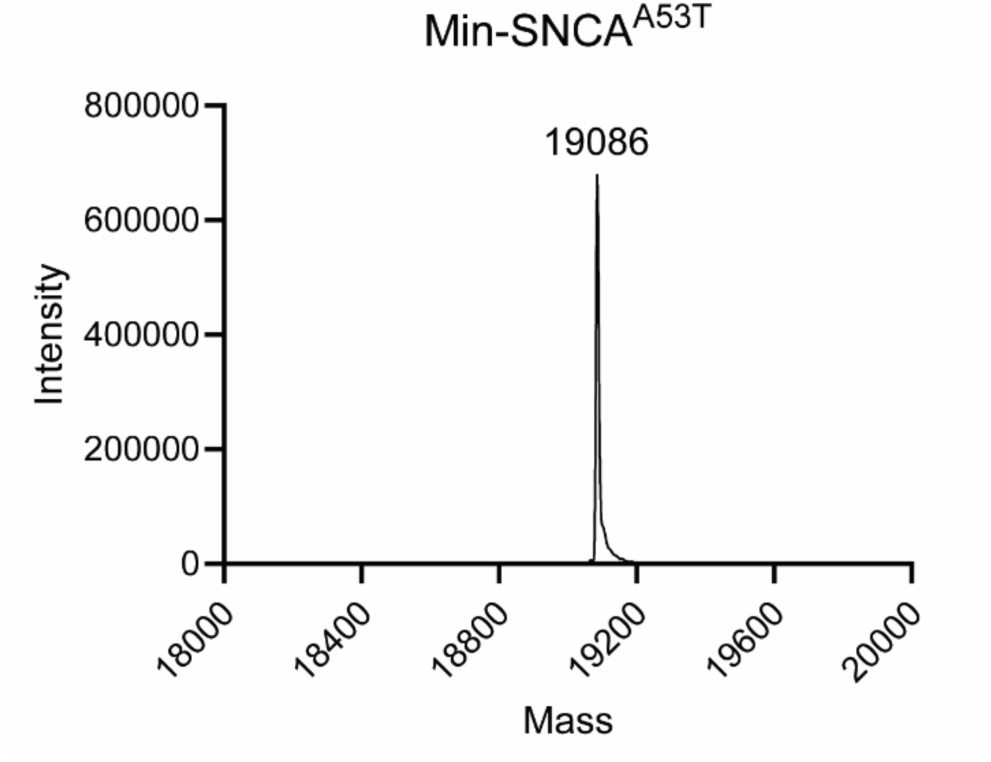
Mass spectrometry analysis of purified degron tagged alpha-synuclein (Min-SNCA^A53T^).

**Supplementary Figure 6.**
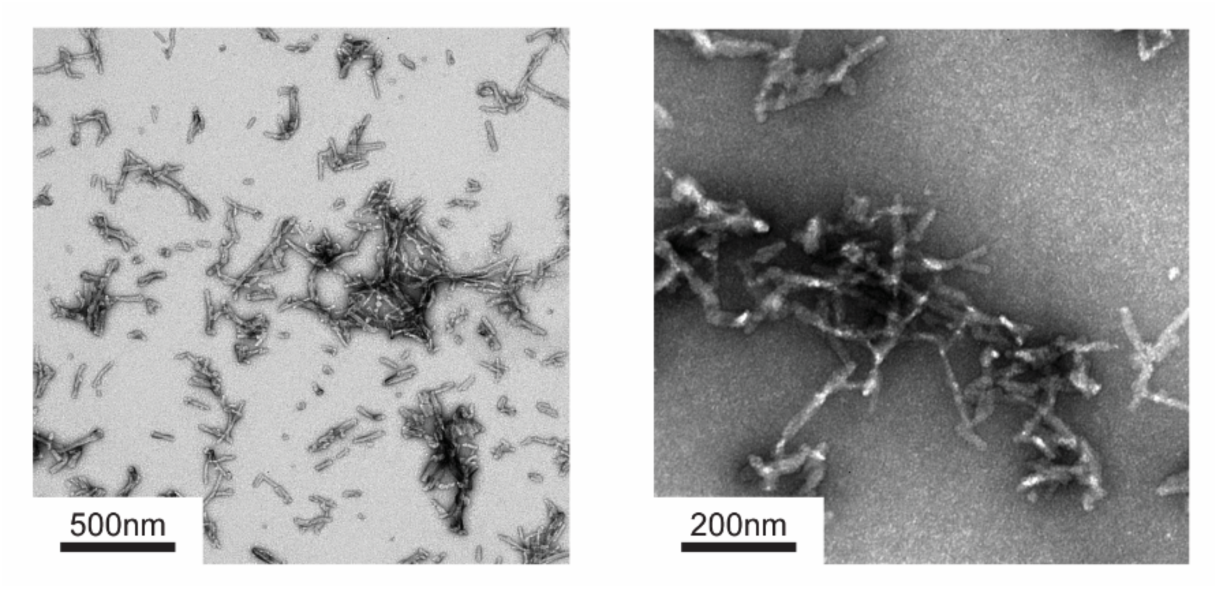
TEM images of co-aggregated alpha-synuclein. (90% degron tag and 10% wild type).

**Supplementary Figure 7.**
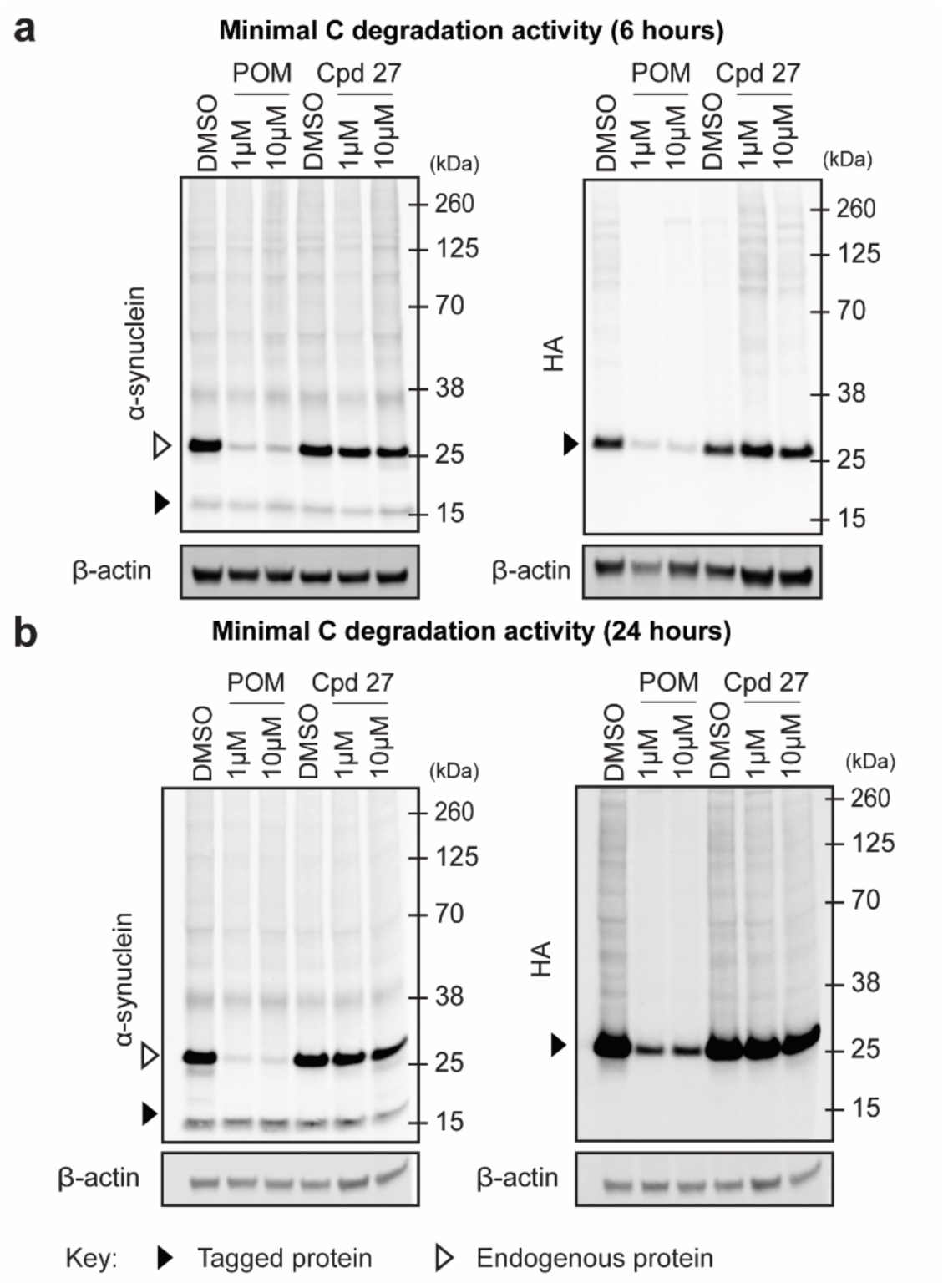
Clerance of preformed fibril aggregated alpha-synuclein using minimal degron tags. **a**. Western blots of alpha-synuclein and HA levels following 6-hour treatment of pomalidomide or compound 27 in SNCA^A53T^-Min expressing cells transfected with PFFs. **b.** Western blots of alpha-synuclein and HA levels following 24-hour treatment of pomalidomide or compound 27 in SNCA^A53T^-Min expressing cells transfected with PFFs.

**Supplementary Figure 8.**
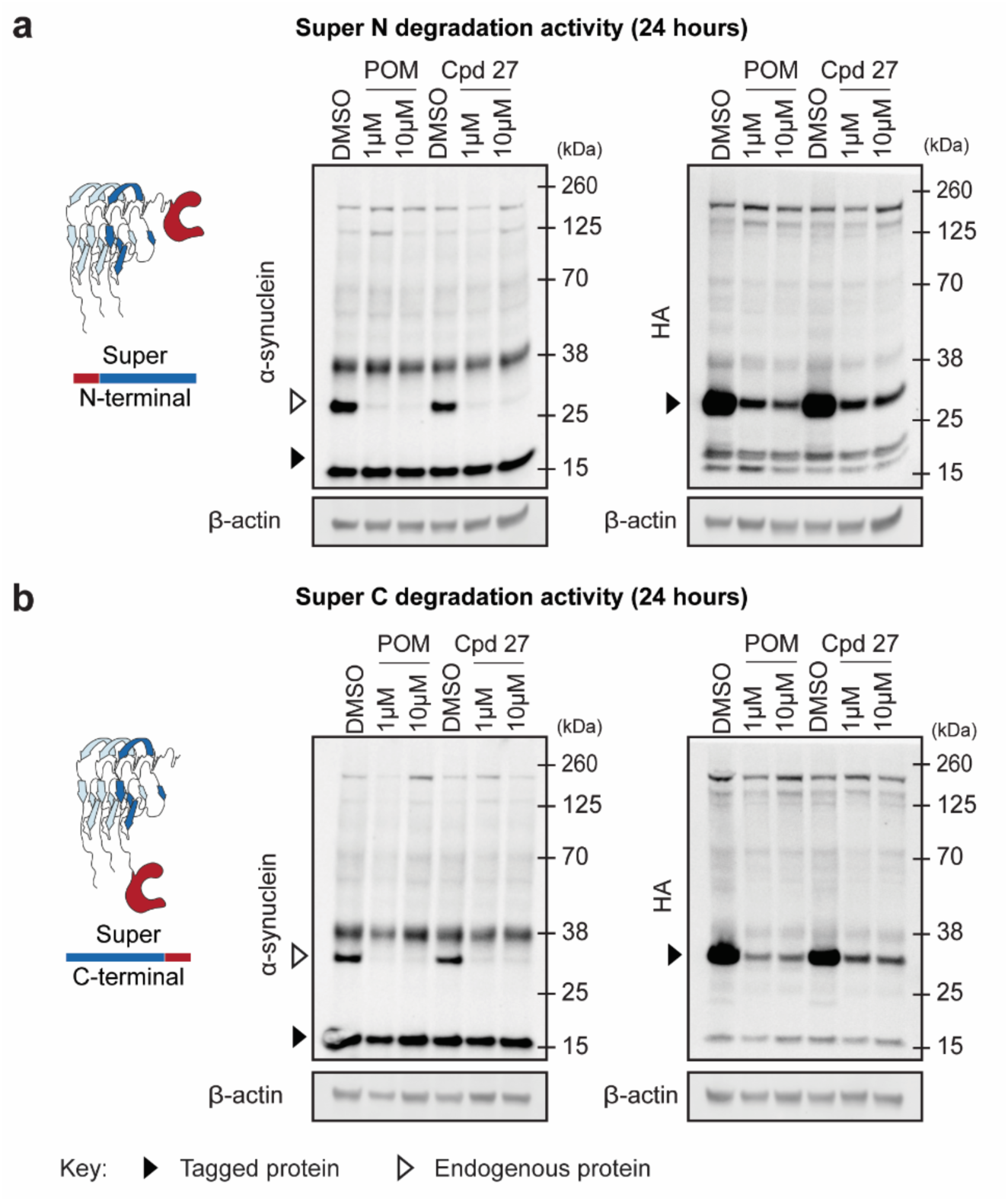
Super degron tagged A53T alpha-synuclein cells transfected with PFF. **a-b.** Western blots of alpha-synuclein and HA levels following treatment of pomalidomide or compound 27 in super degron tagged A53T alpha-synuclein transfected with PFF. HA tag on degron-tagged alpha-synuclein. **a.** N-terminal tagged **b.** C-terminal tagged.

**Supplementary Figure 9.**
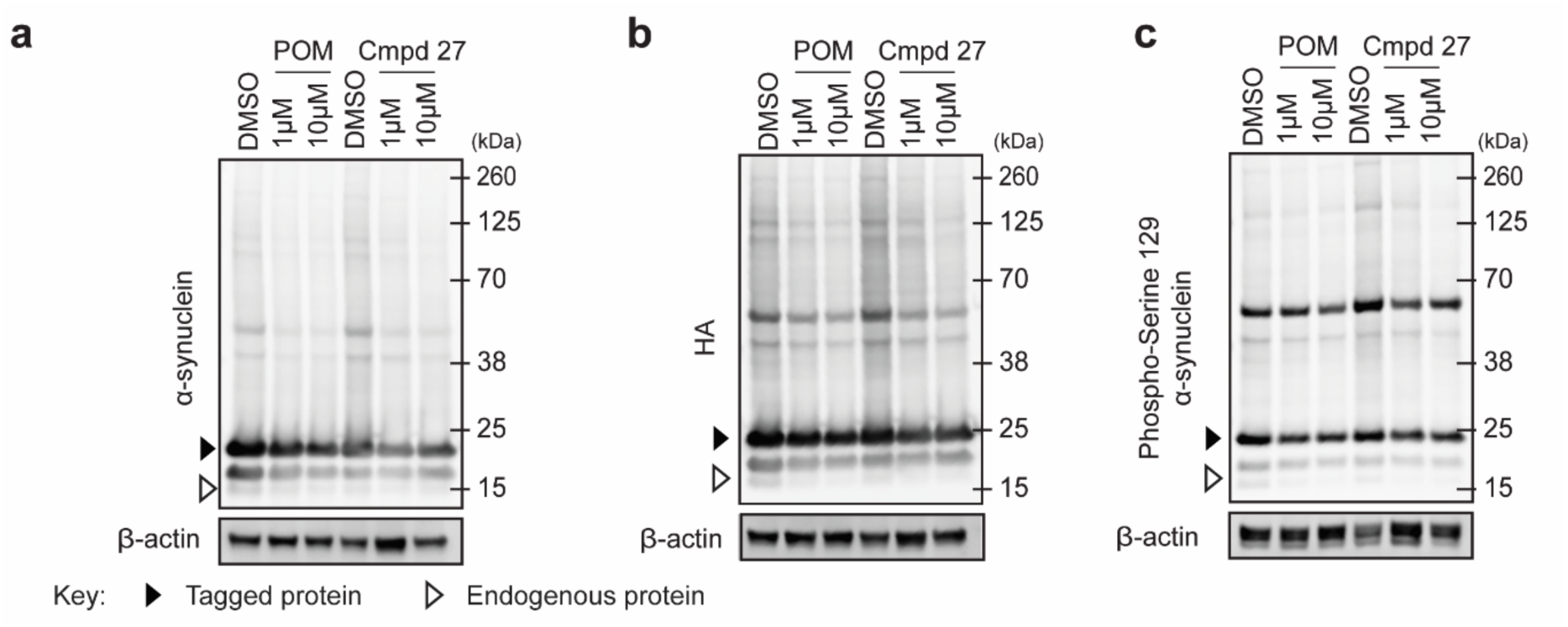
Western blots of SHSY5Y neurons expressing Min-SNCA^A53T^ via CMV promoter treated with PFFs for 24 hrs, followed by 24-hour compound treatment. **a.** Total alpha-synuclein. **b.** Degron tagged alpha-synuclein. **c.** Disease relevant pS129 oligomers.

