## Supplementary material for "A Degron Decoy System Co-opts Pathological Seeding to Enable Clearance of Multimeric α-Synuclein": Uncropped Blots

**Figure 1C**

6 hours

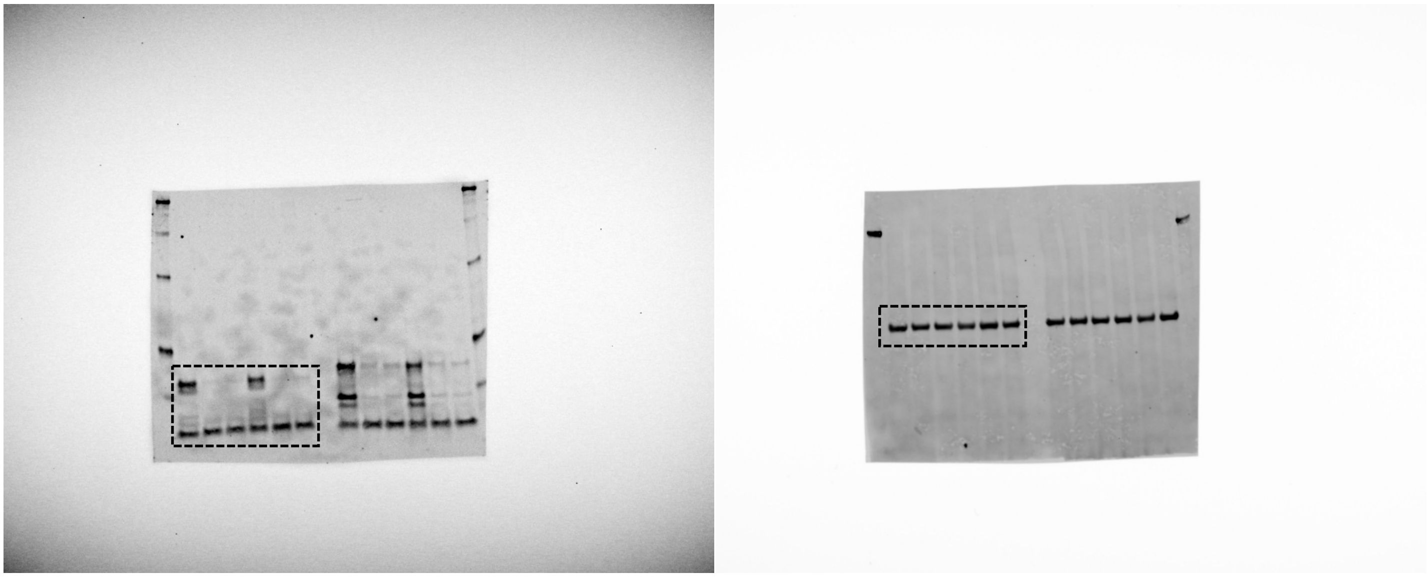

24 hours

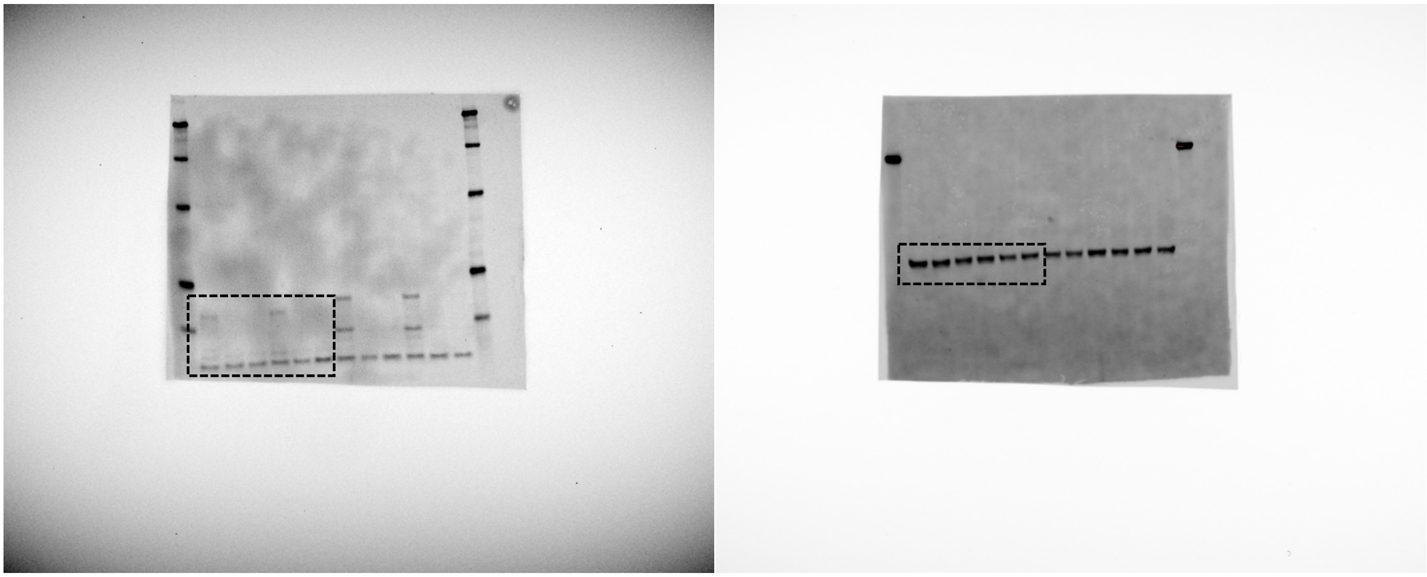

**Figure 1D**

6 hours

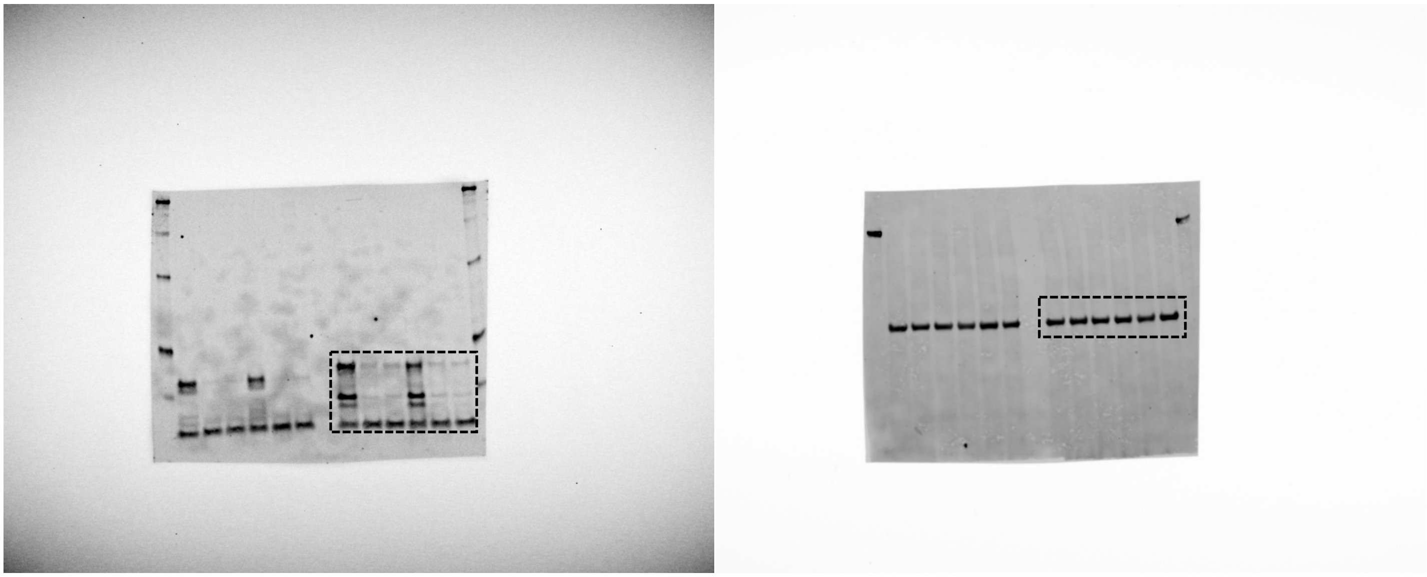

24 hours

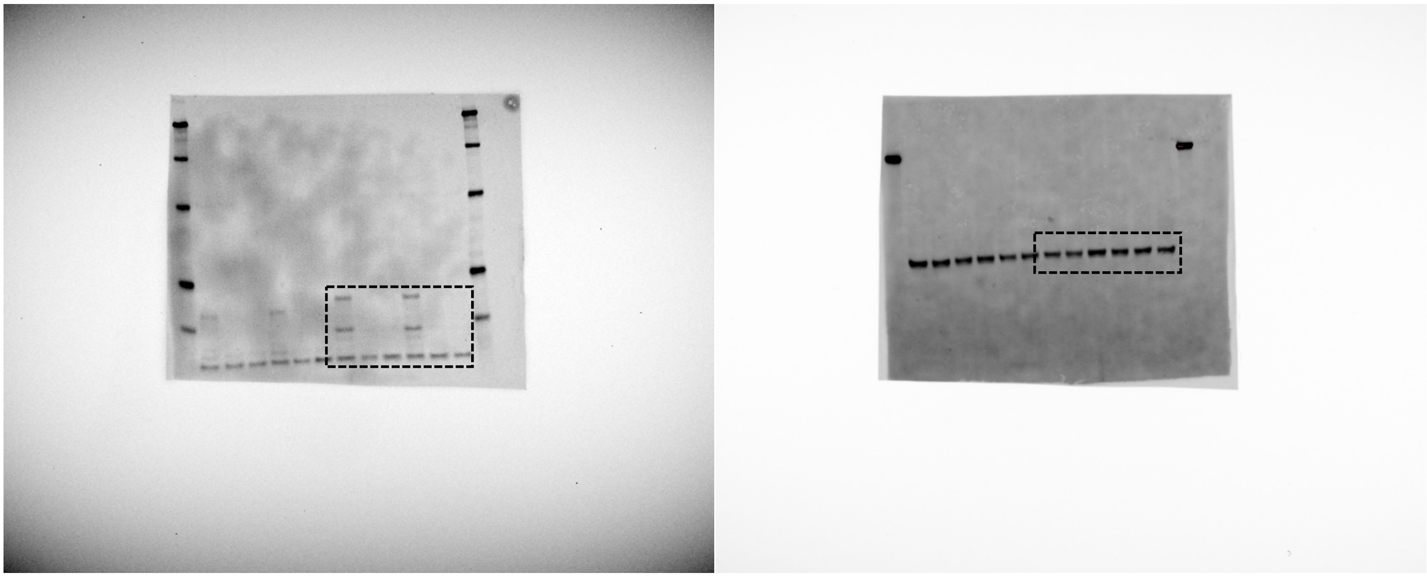

**Figure 1E**

6 hours

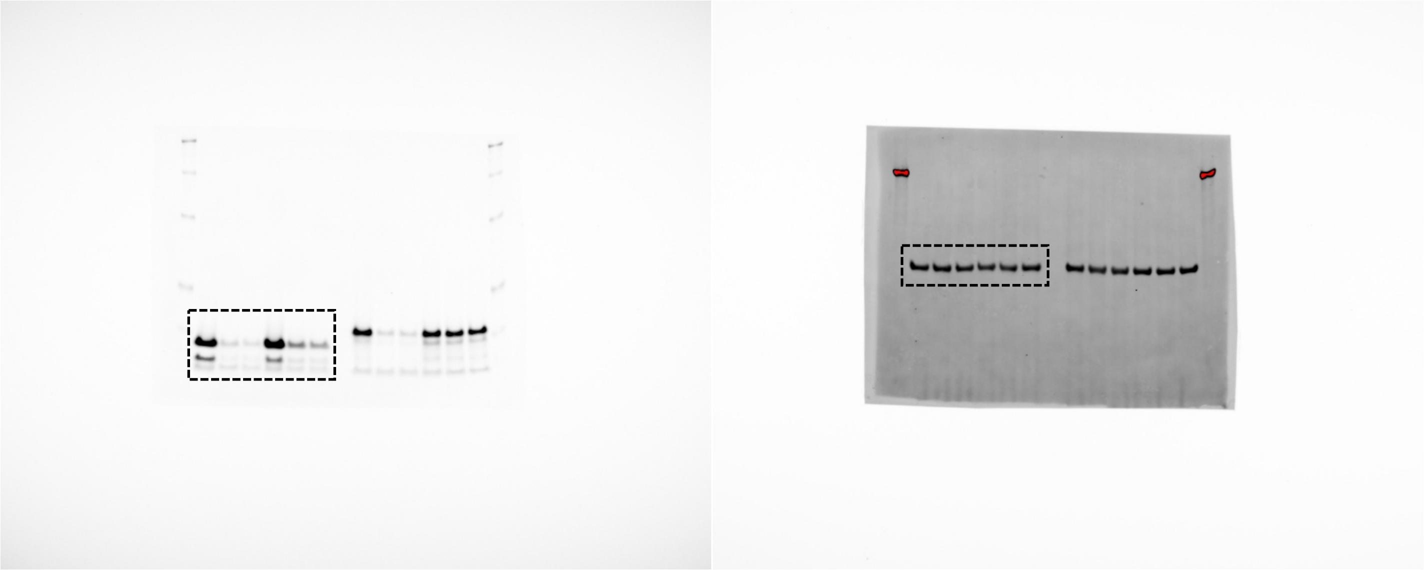

24 hours

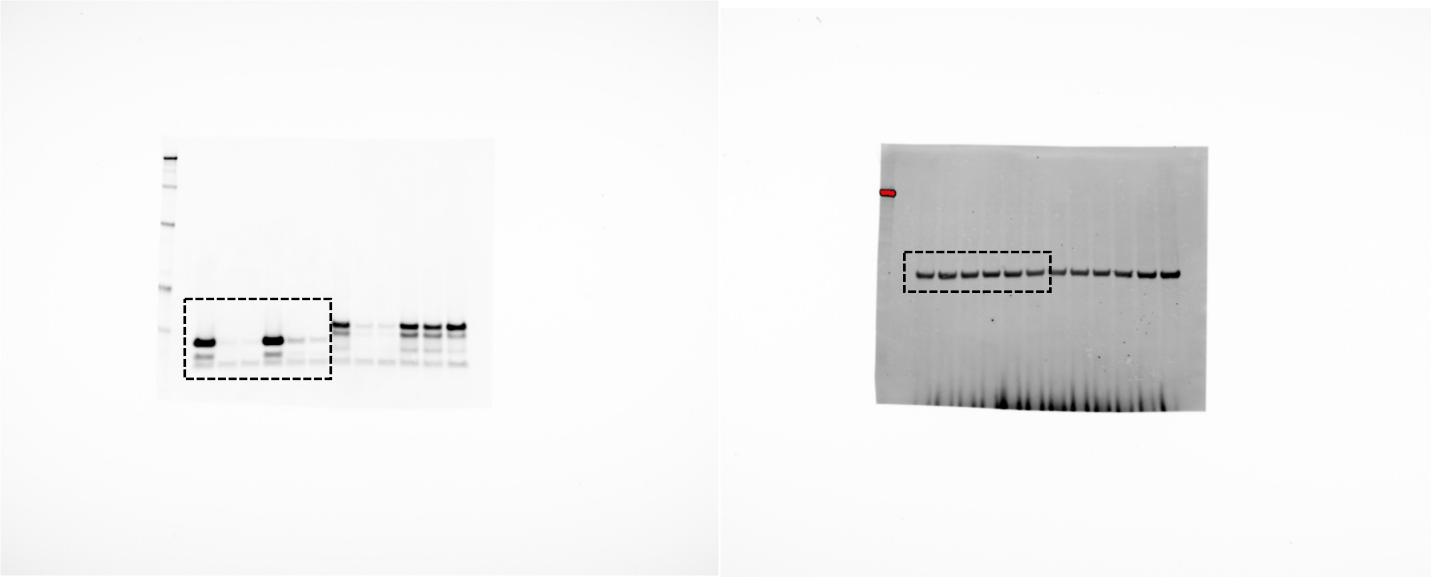

**Figure 1F**

6 hours

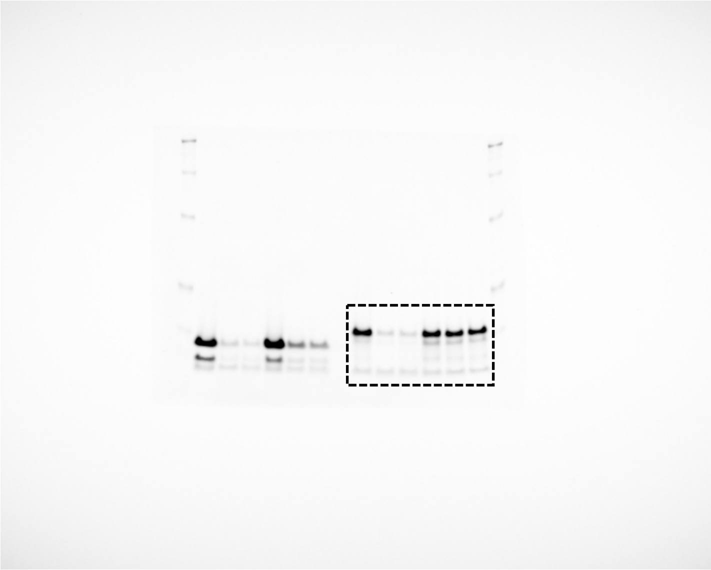

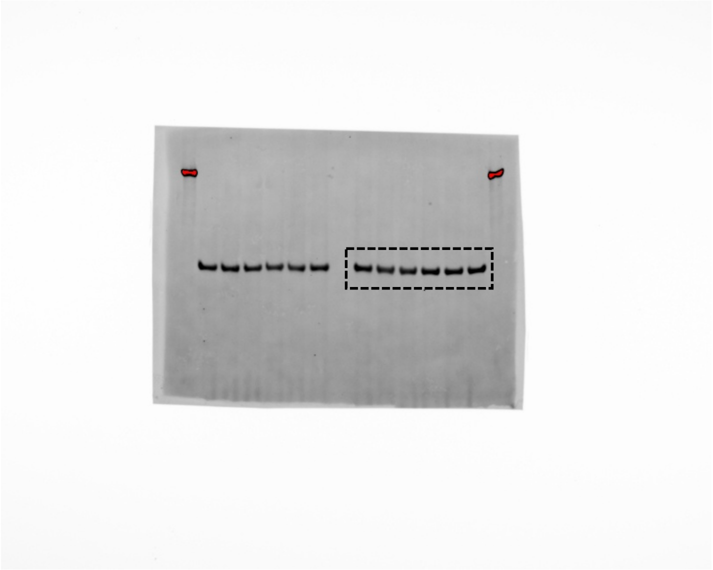

24 hours

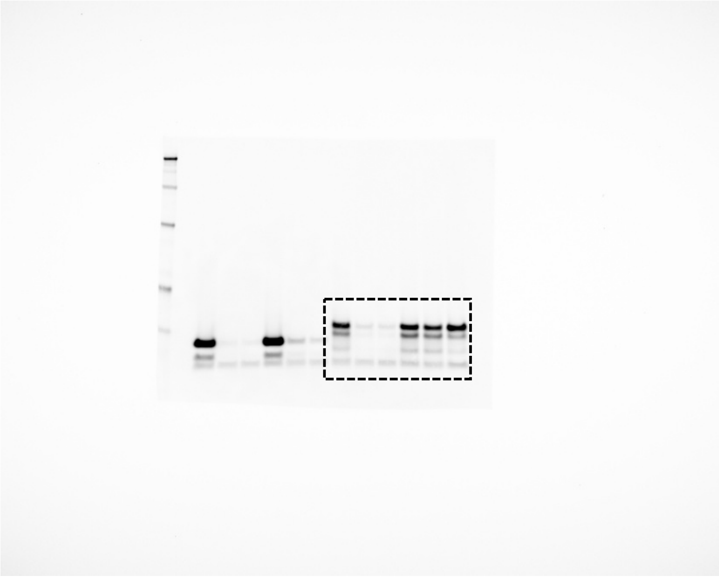

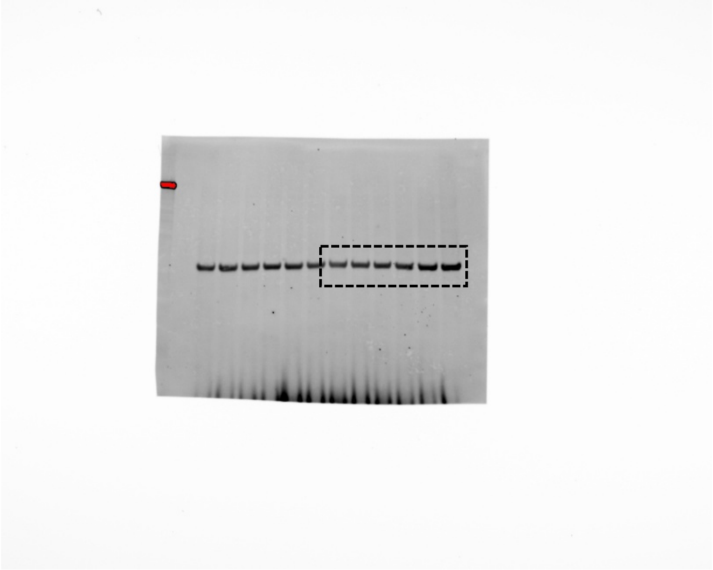

**Figure 3B**

Alpha syn

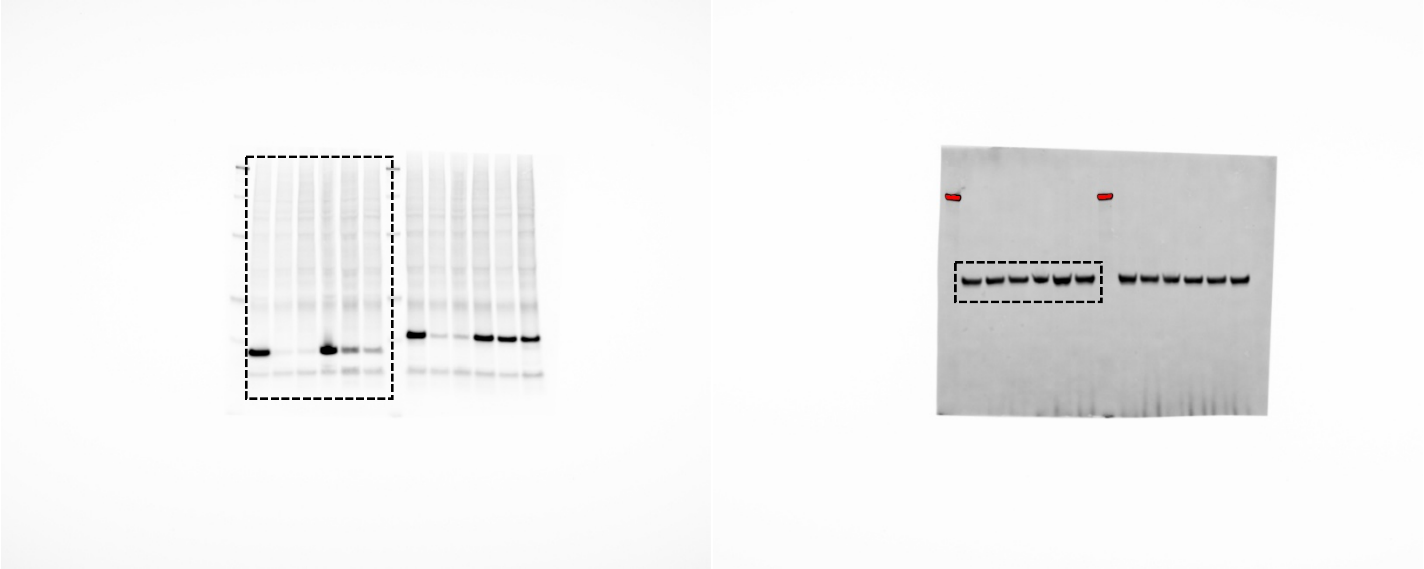

HA

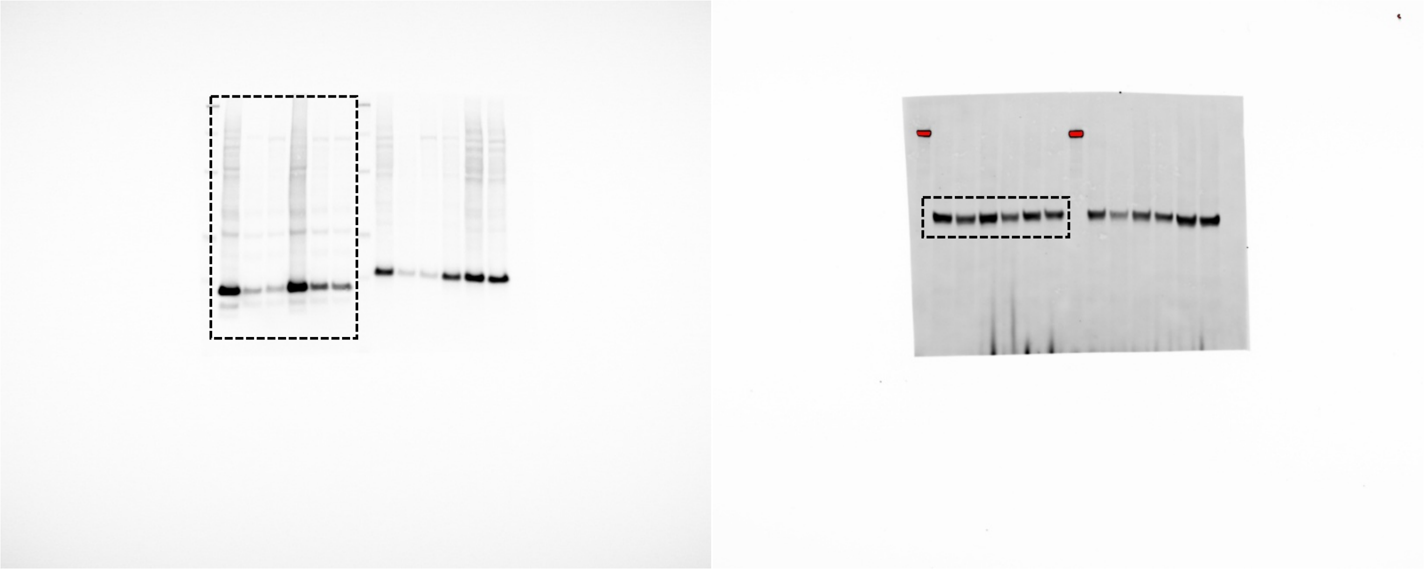

**Figure 3D**

Alpha-synuclein

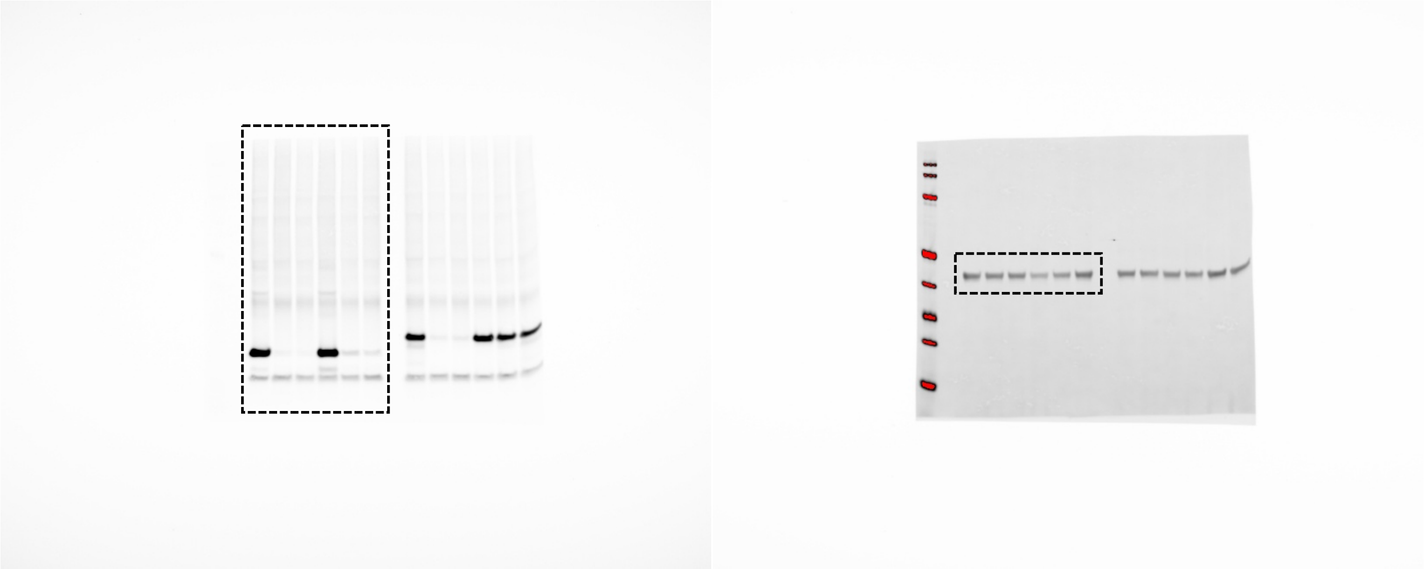

HA

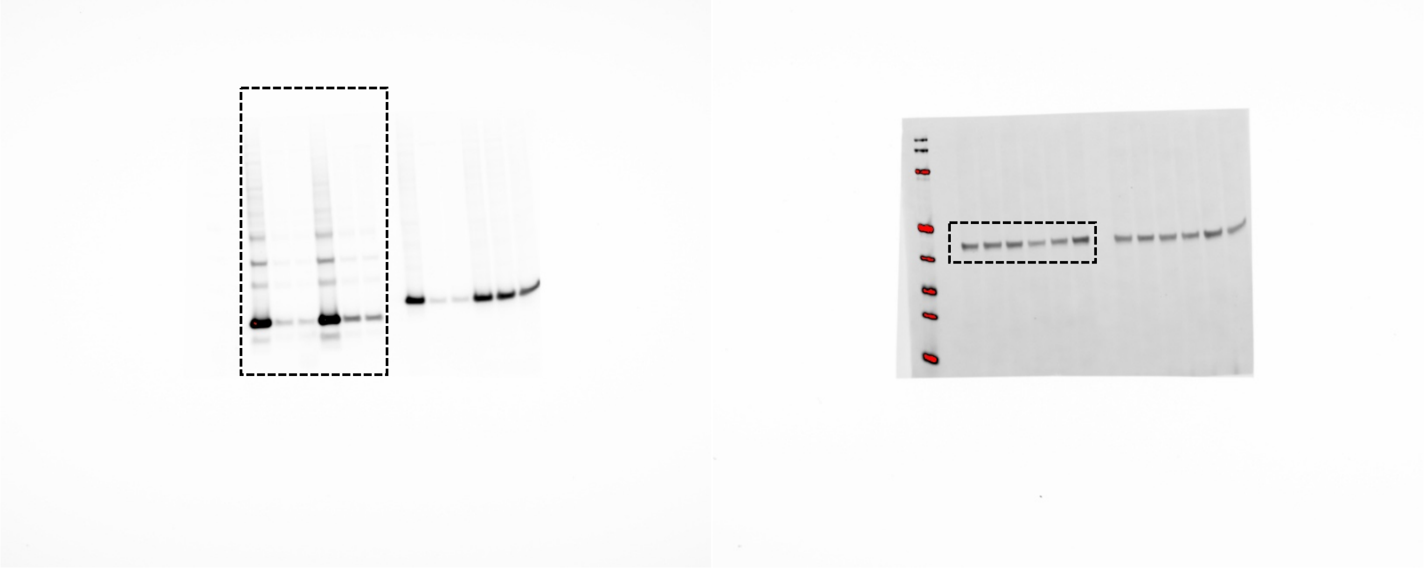

**Figure 4A**

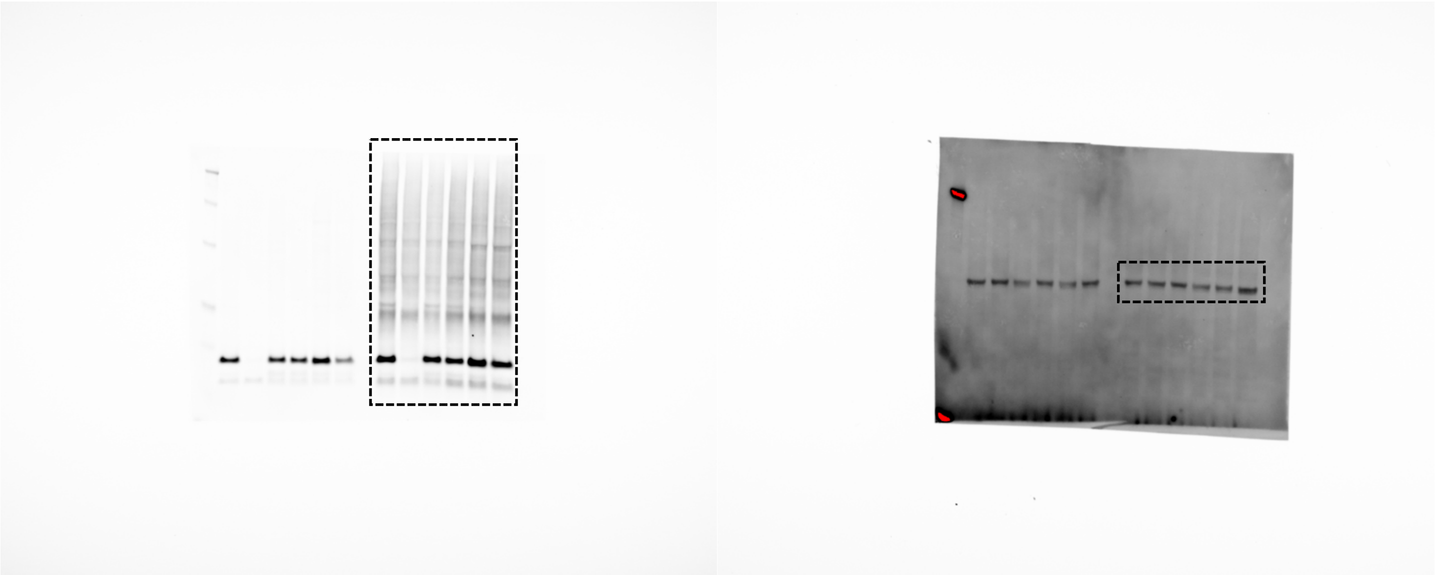

**Figure 4B**

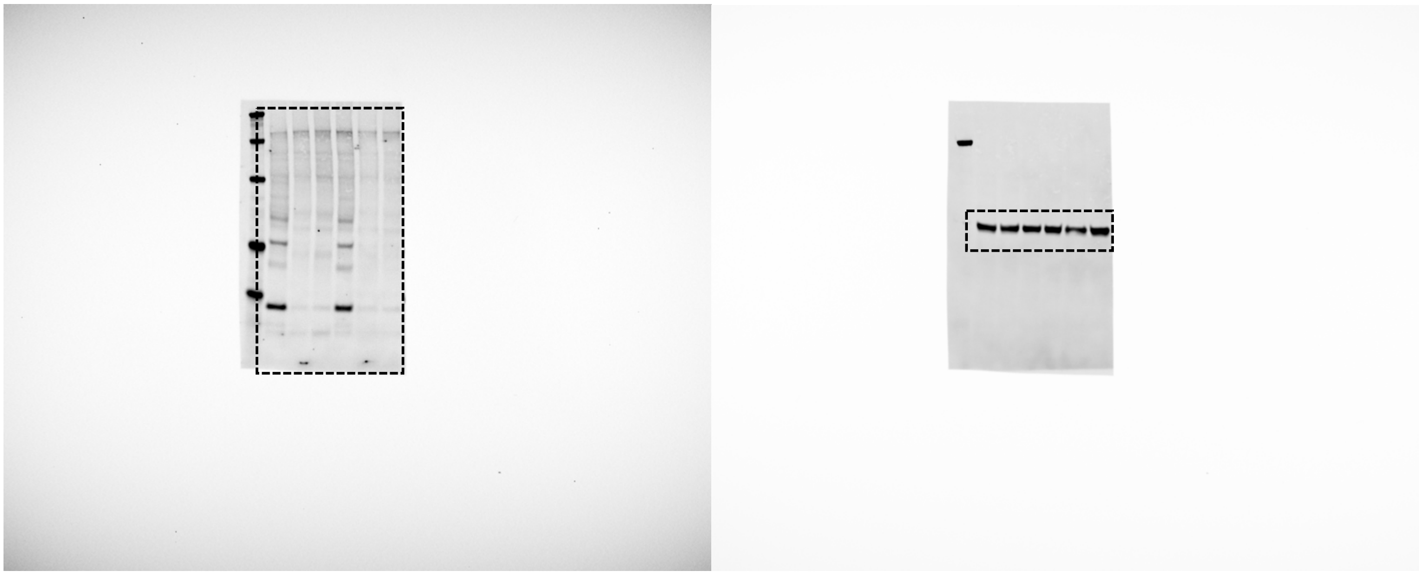

**Figure 4C**

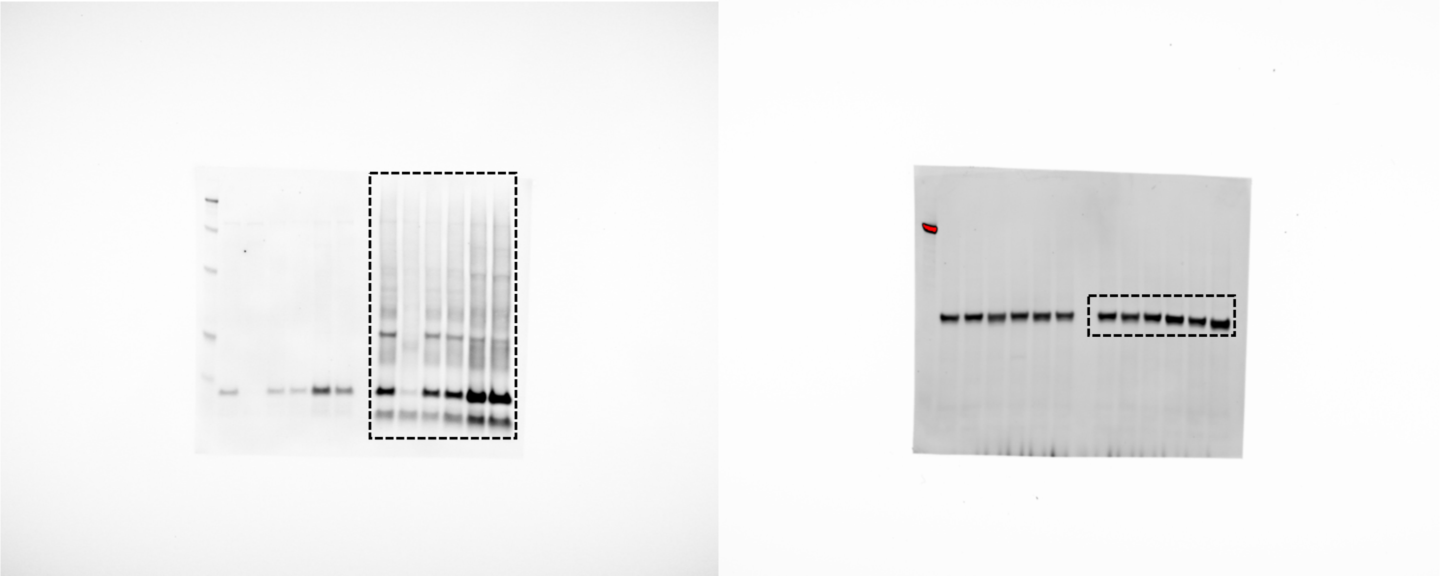

**Figure 5B**

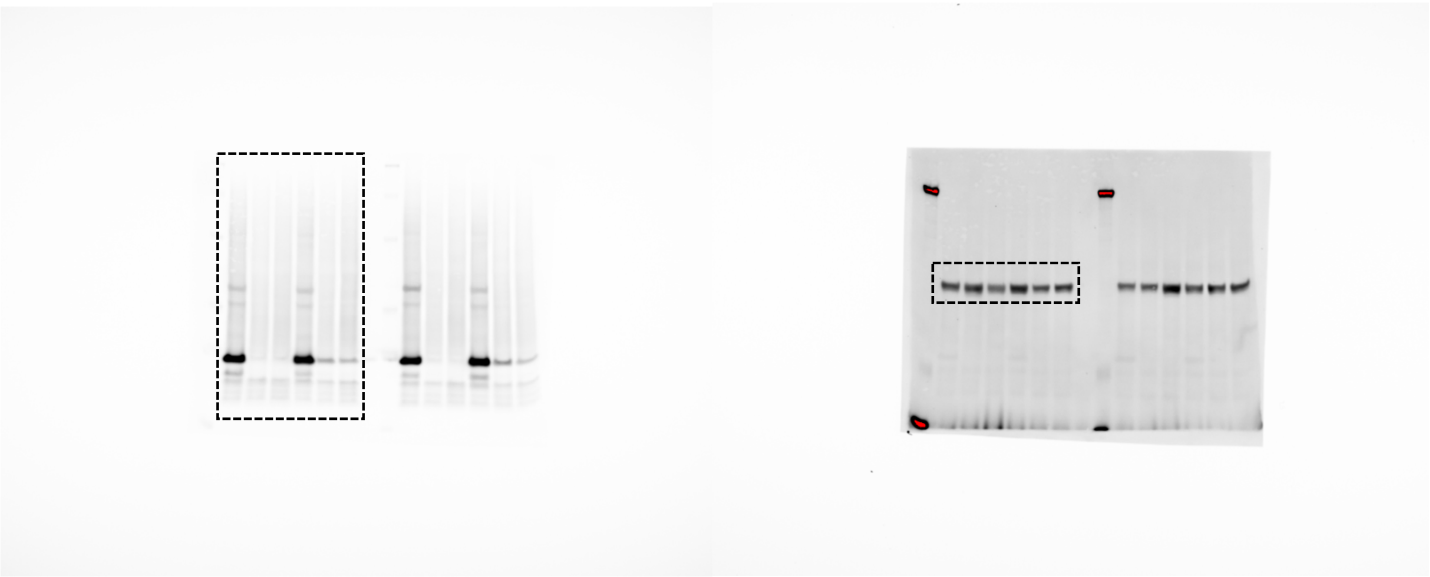

**Figure 5C**

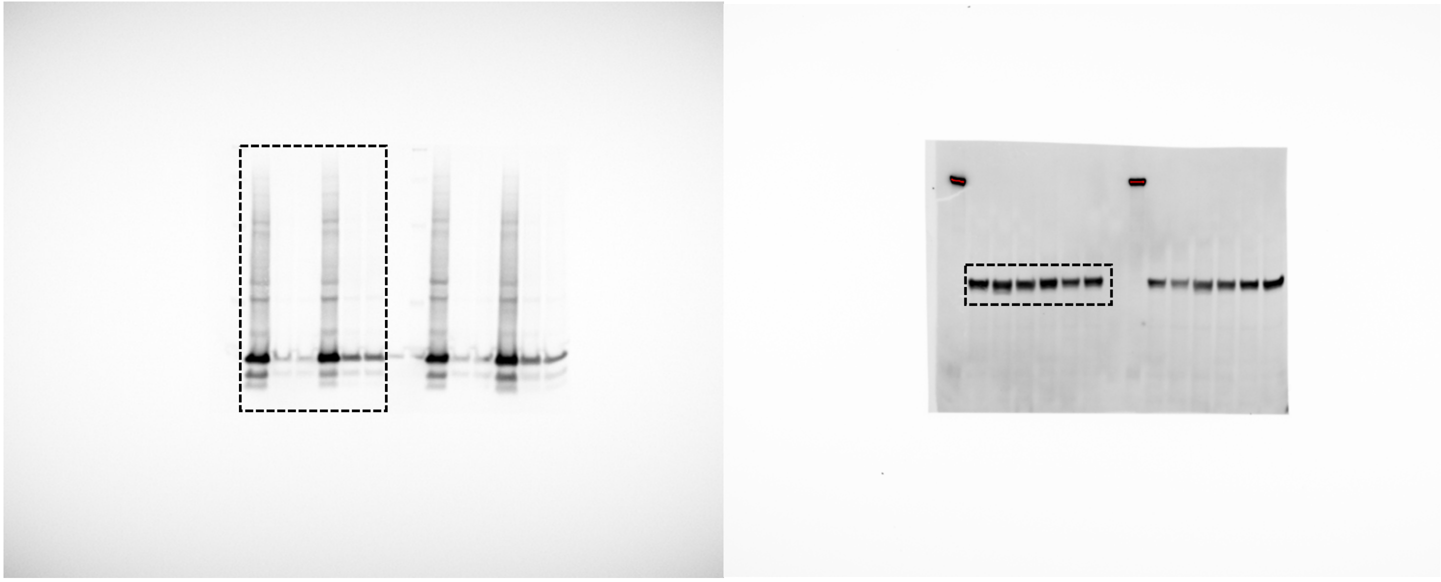

**Figure 5D**

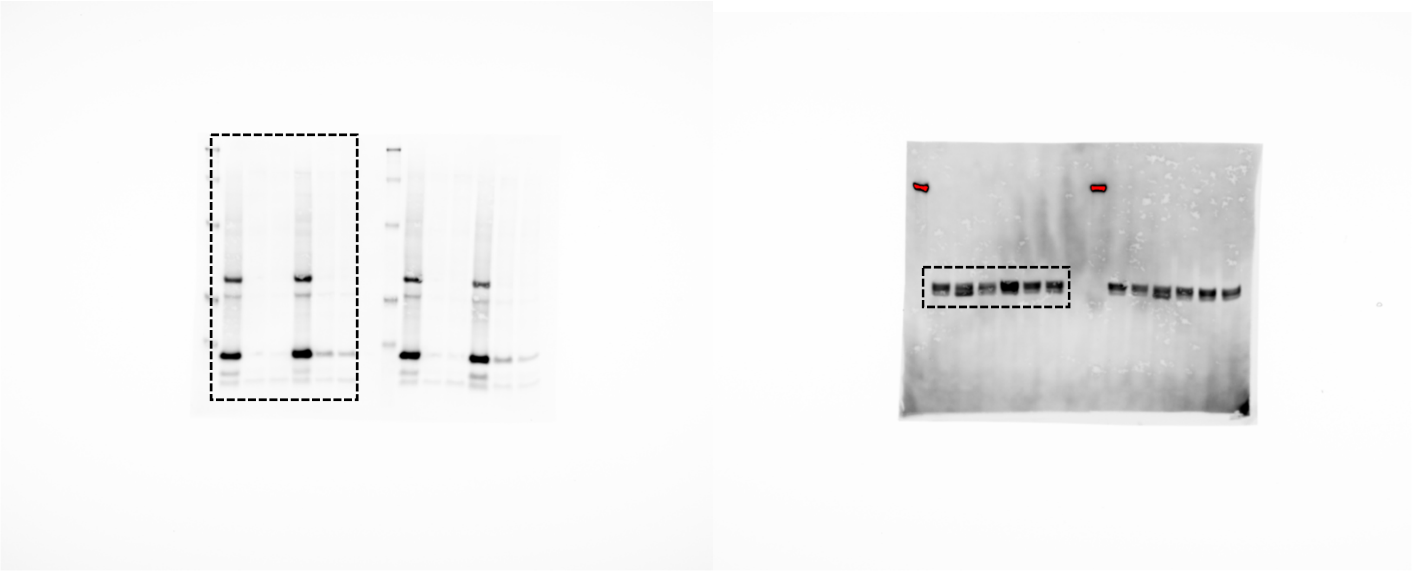

Figure 5E

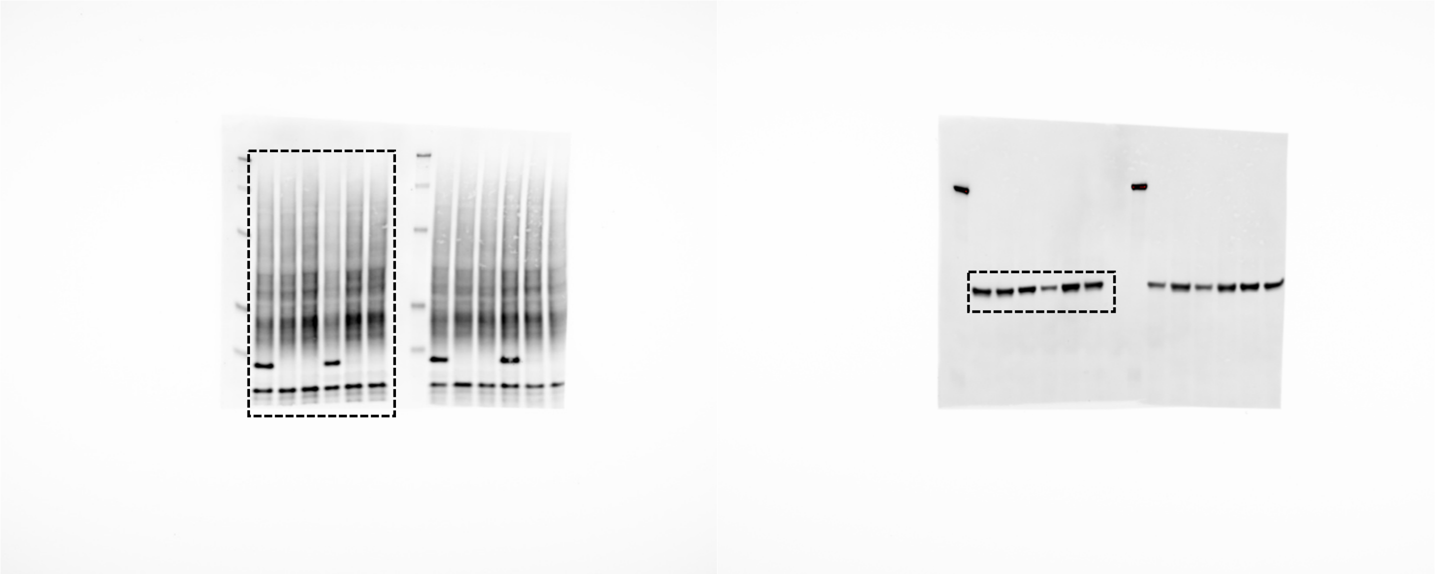

Figure 5F

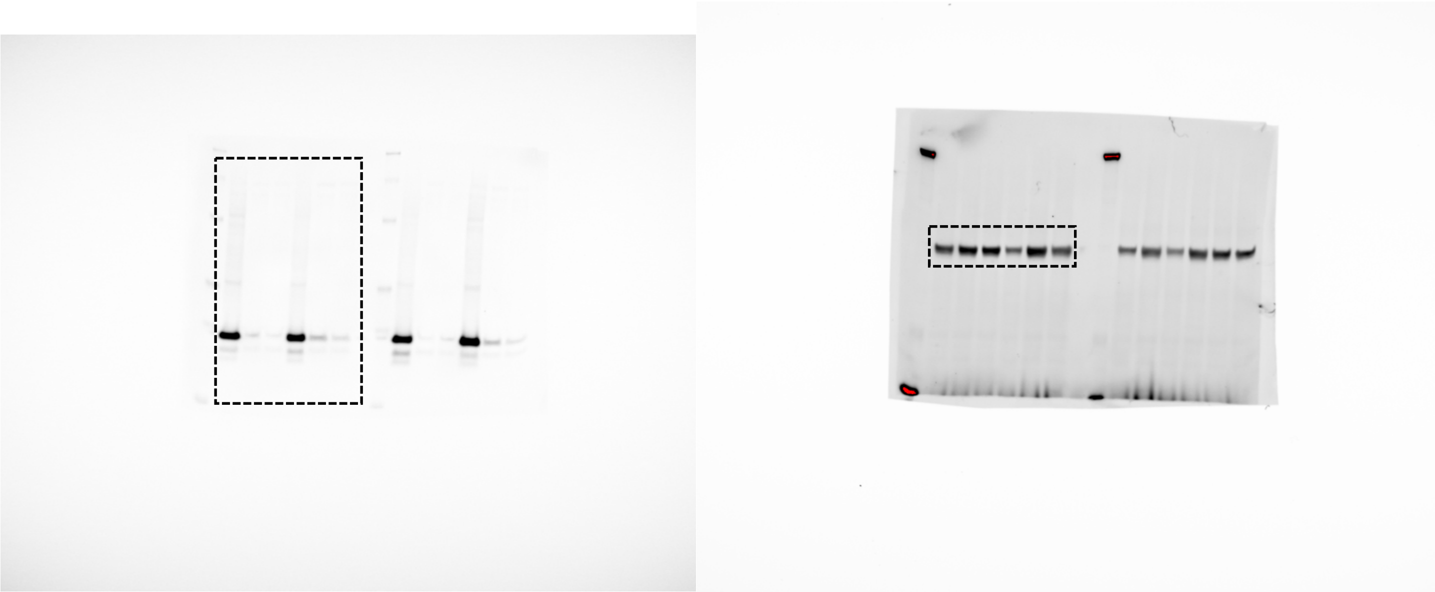

**Figure 5G**

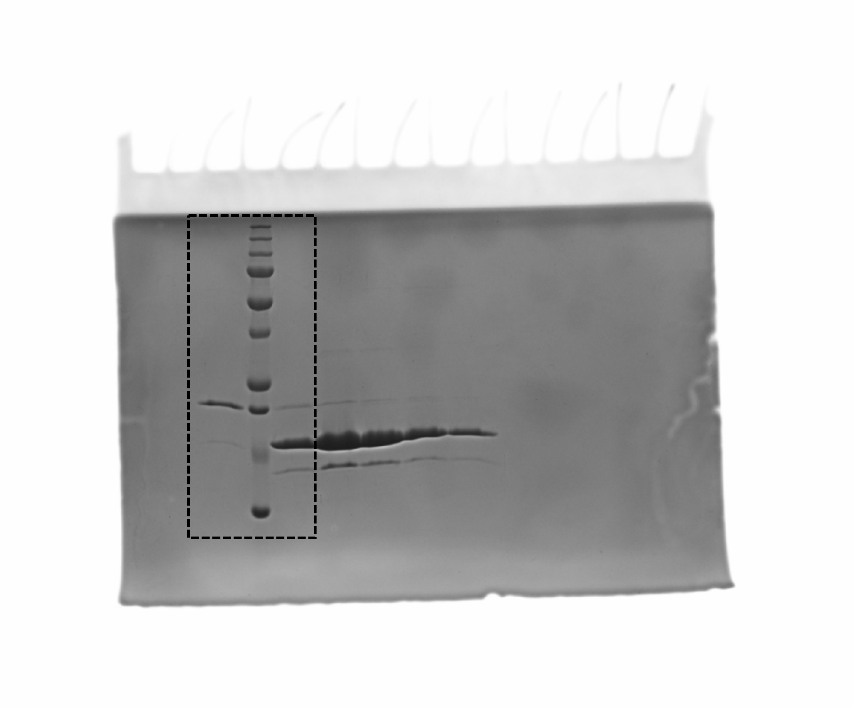

**Supplemental Figure 3A**

Coomassie blue

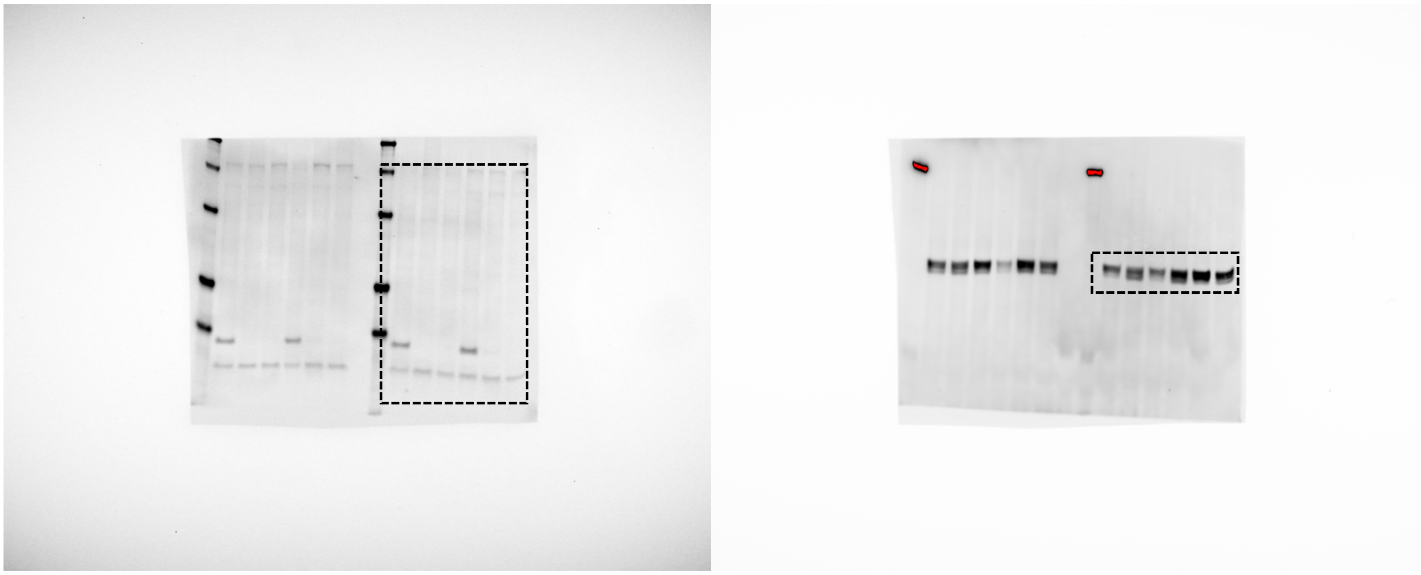

**Supplemental Figure 7A**

Alpha-synuclein

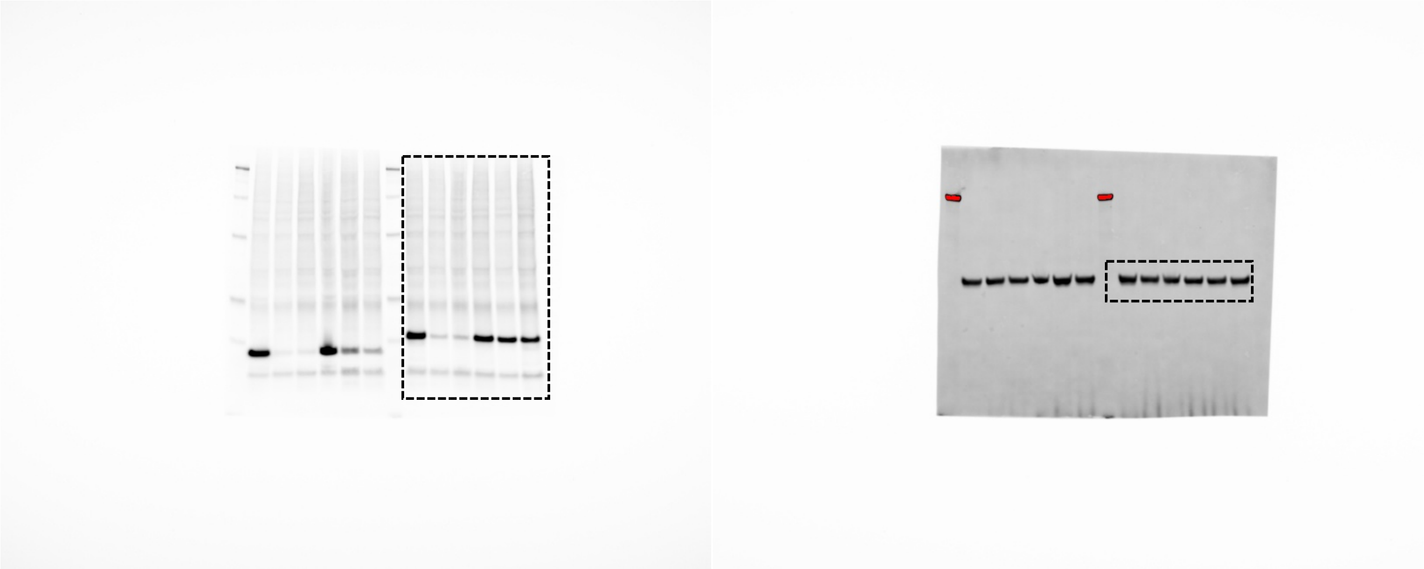

HA

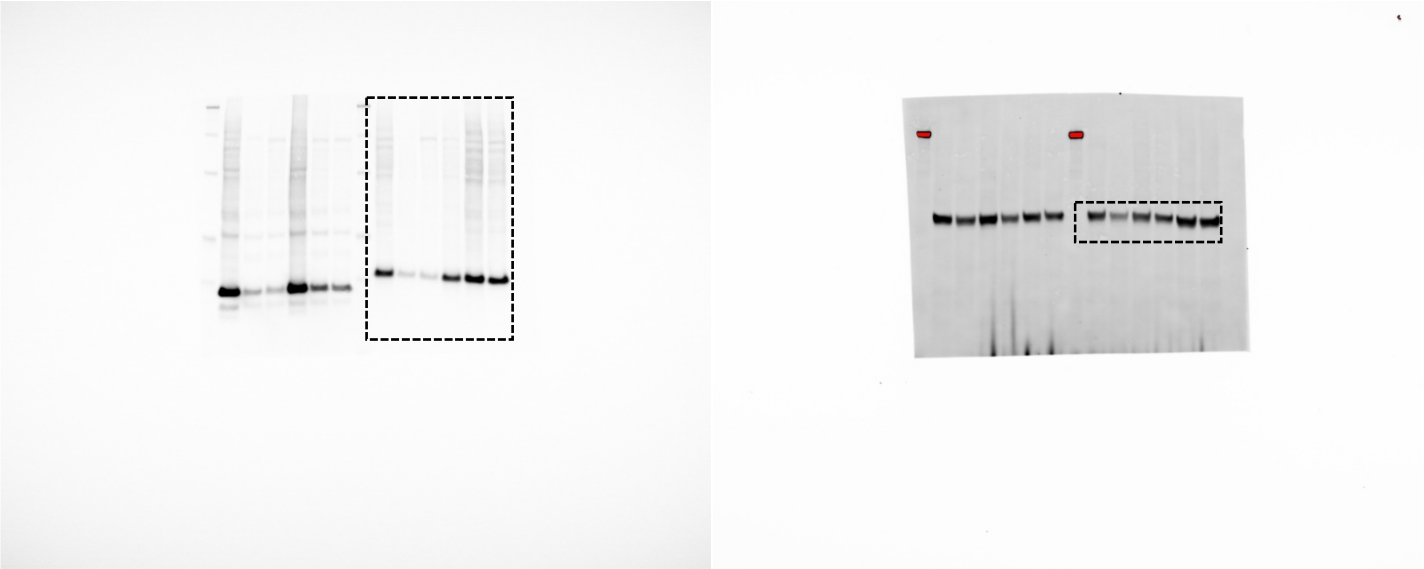

**Supplemental Figure 7B**

Alpha synuclein

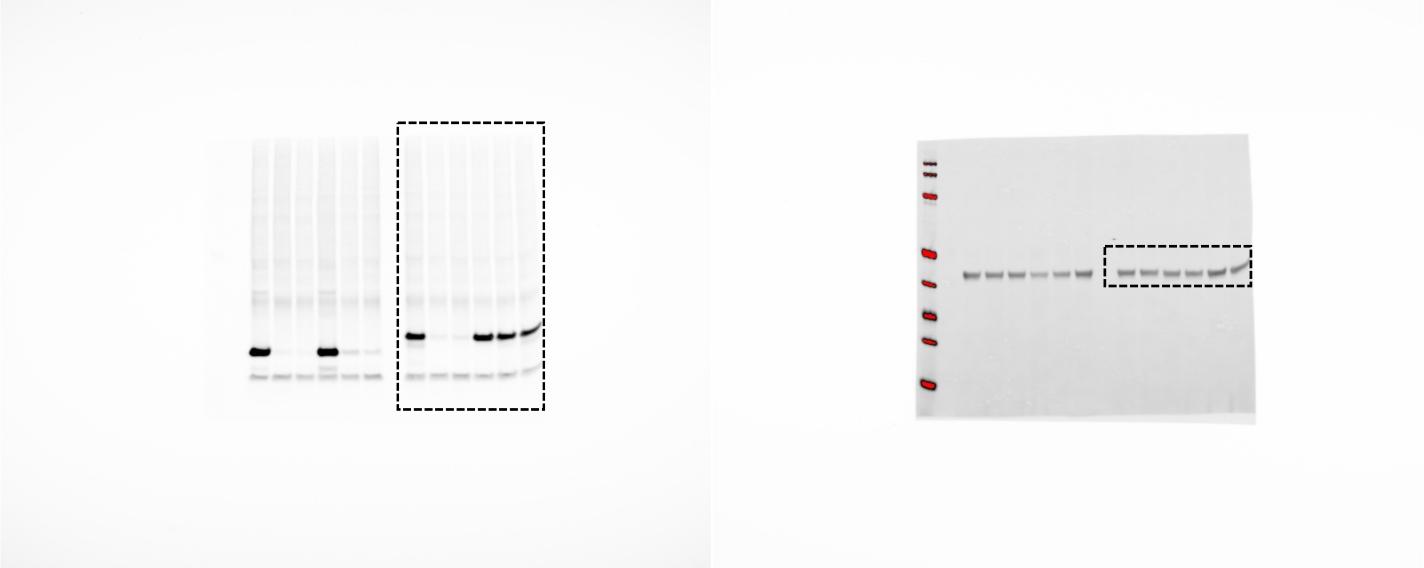

HA

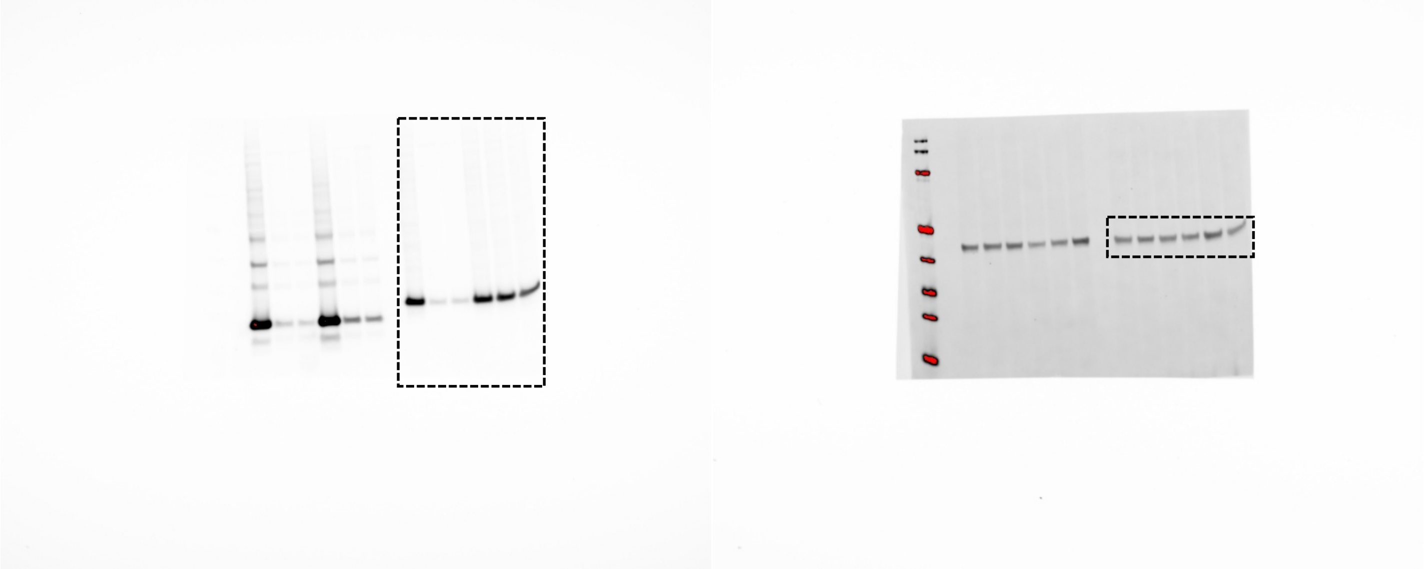

**Supplemental Figure 8A**

Alpha-synuclein

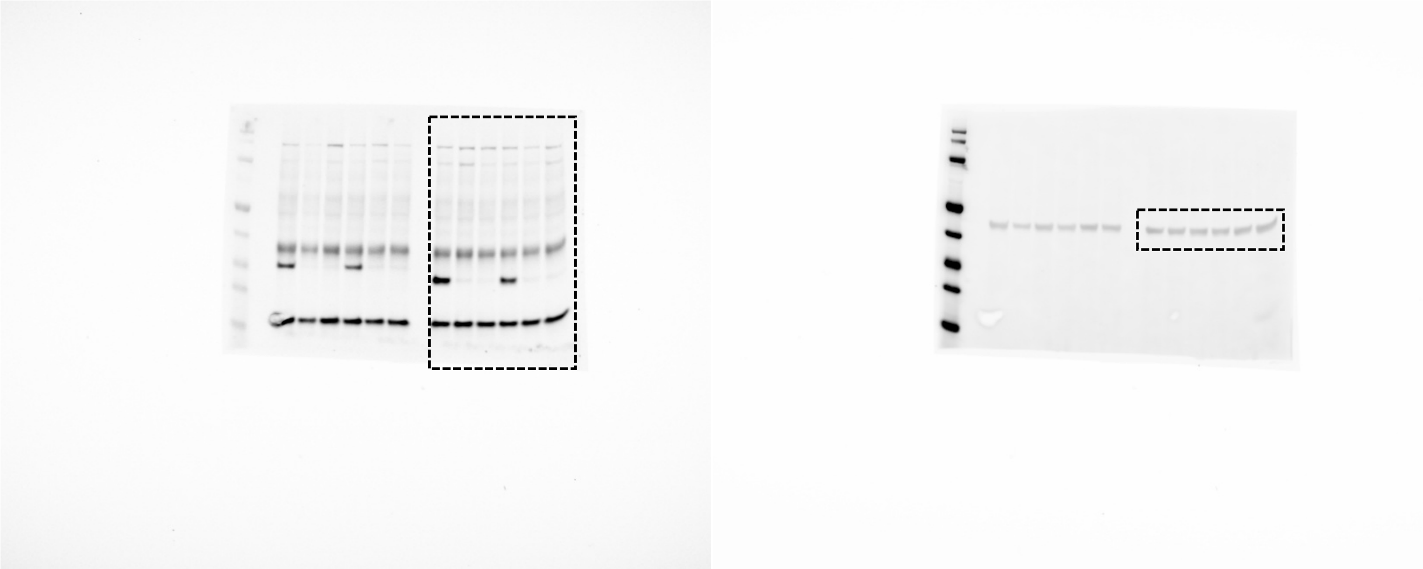

HA

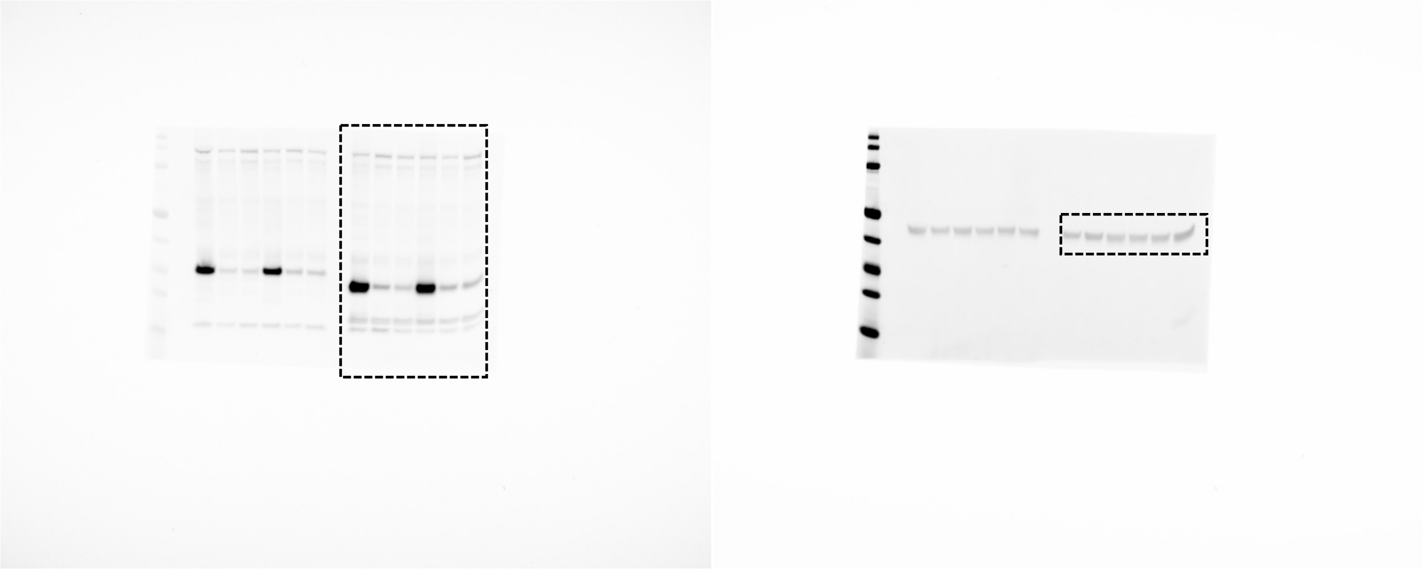

**Supplemental Figure 8B**

Alpha-synuclein

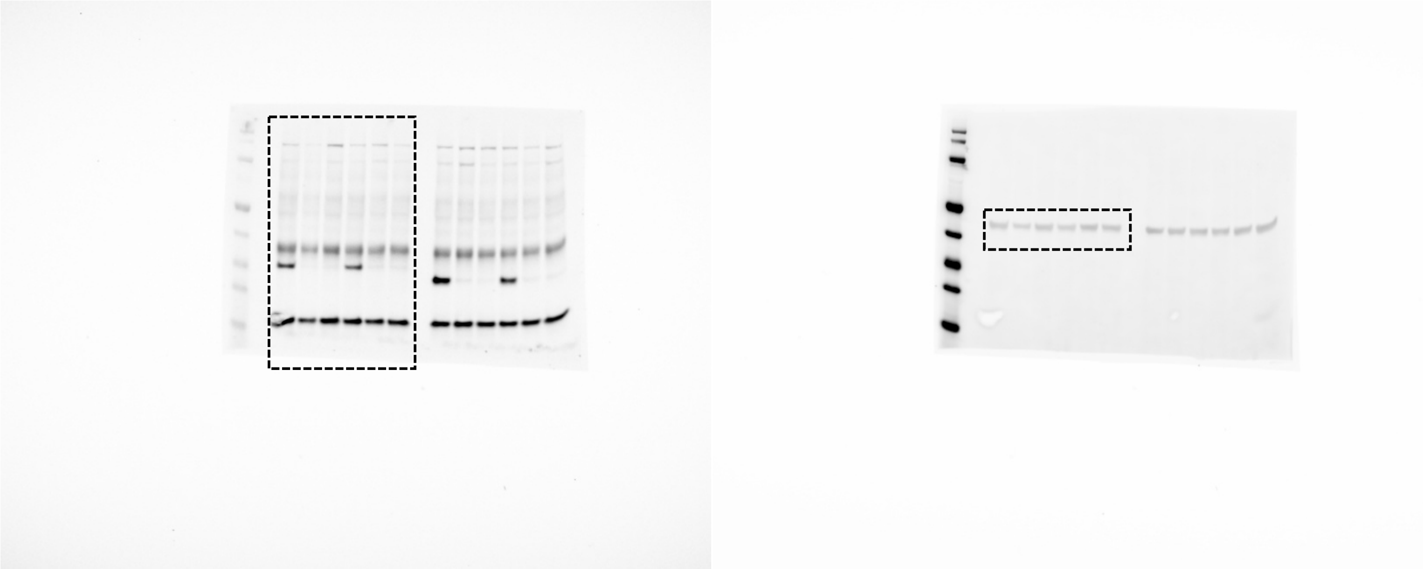

HA

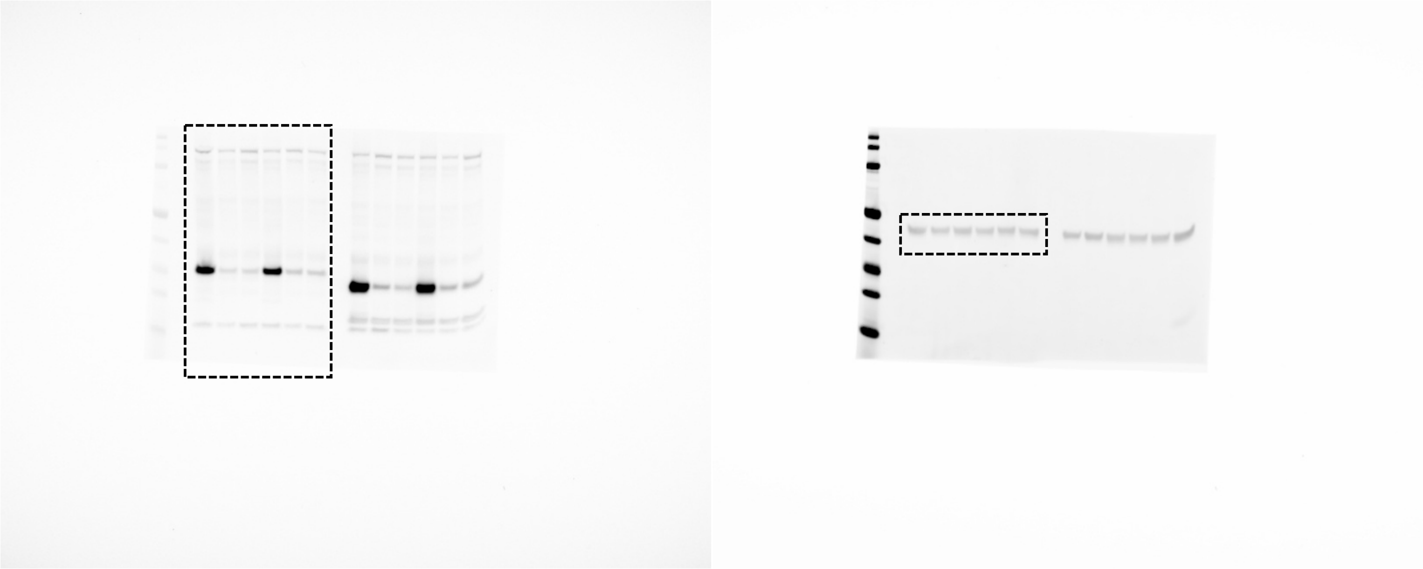

**Supplemental Figure 9A**

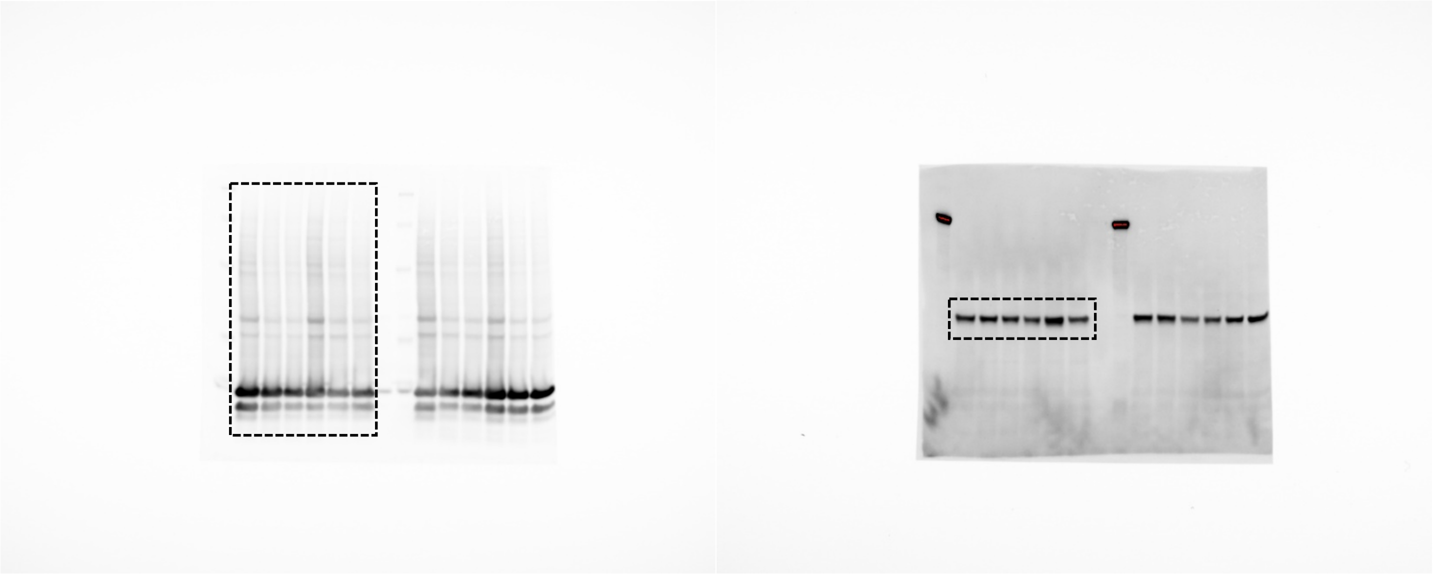

**Supplemental Figure 9B**

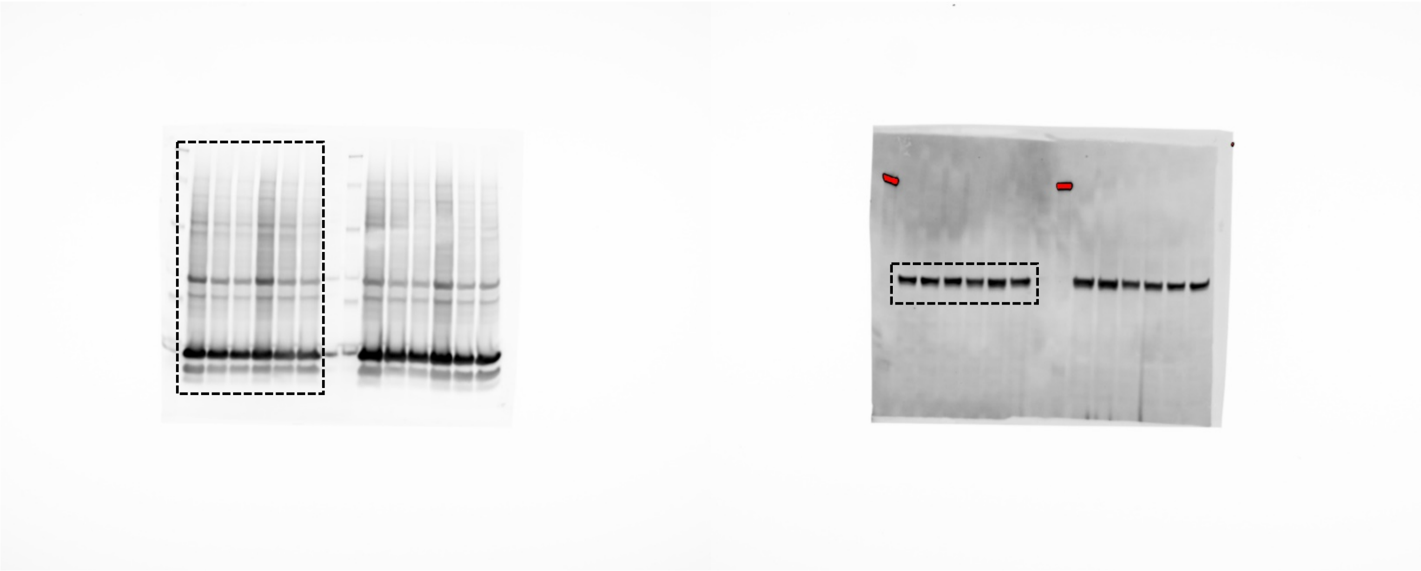

**Supplemental Figure 9C**

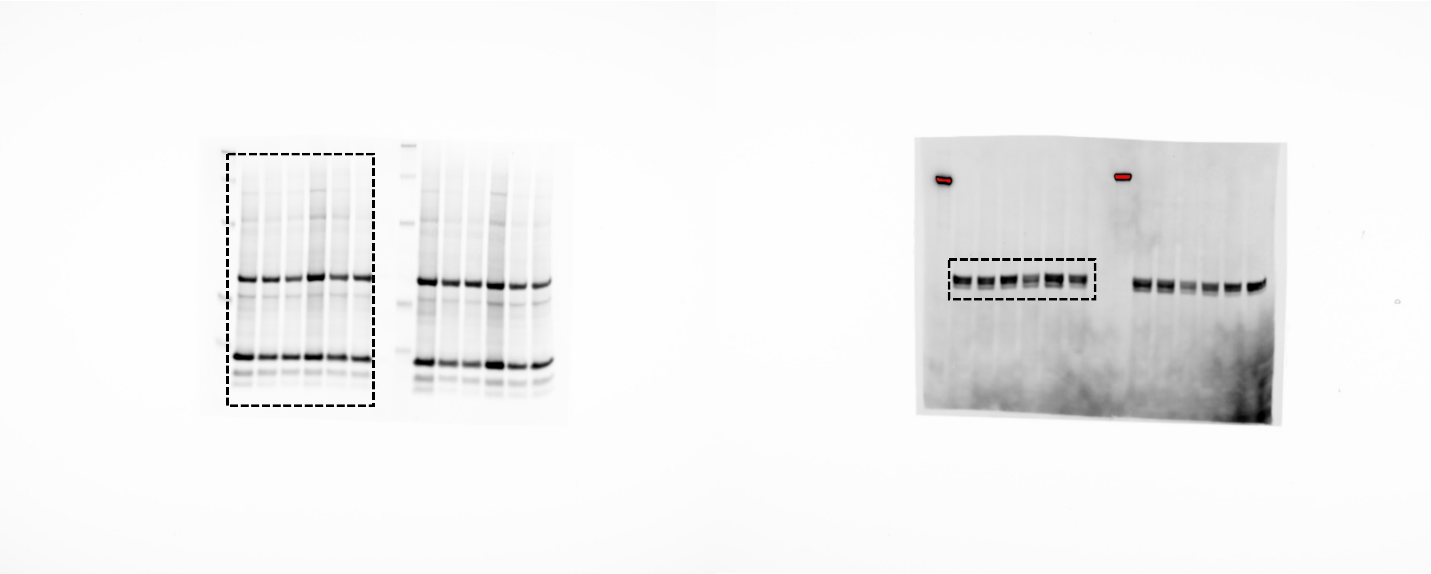
