## Supplementary material for "A Degron Decoy System Co-opts Pathological Seeding to Enable Clearance of Multimeric α-Synuclein": SI Methods

**Cell culture**
The following cell lines were employed in this study: HEK293 (ATCC, CRL-1573) and SHSY5Y cells (ATCC, CRL-2266) were cultured in high-glucose DMEM (Gibco, 11965092) with 10% FBS (Gibco, 10437028) and 1% Penicillin–Streptomycin (Gibco, 15140122). were cultured in All cell lines were maintained in 37 °C and 5% CO2 incubators and routinely tested negative for mycoplasma contamination using the MycoAlert Kit (Lonza, LT07318).

**Plasmids**
Custom plasmids containing degron tag A53T alpha-synuclein constructs were cloned into pcDNA3.1(+) backbone by Genewiz from Azenta Life Sciences. Lentiviral plasmids containing degron tag A53T alpha-synuclein were cloned into pTwist Lenti CAG Puro or pTwist Lenti TRE Puro by Twist Biosciences.

**Cell line generation**
HEK293 cells were plated into a 6-well plate (Corning, 3513) at 800,000 cells per well in culture media. Cells were returned to 37 °C and 5% $CO_2$ incubator to adhere to plates overnight. The following day, cells were transfected with 2 μg of plasmid using TransIT-2020 transfection reagent (Mirus, 5404). The cells were selected with 600 μg/mL G418 antibiotic (Gibco, 10131035) for stable expression of constructs. For the SHSY5Y lentiviral lines, lentivirus was produced by Sanford Burnham Prebys' Viral Vector core facility. The SHSY5Y cells were transduced following the BROAD Spinfection protocol using 4 μg/mL polybrene (Sigma, H9268). The cells were selected with 1 μg/mL puromycin (Fisherbrand, BP2956100) for stable expression of the construct. For the cell line with the TRE promoter, 1 μg/mL doxycycline (Thermo Scientific, 446060050) was added 24 hours prior to addition of fibrils.

**SHSY5Y neuron differentiation**
SHSY5Y cells were plated into a 6-well plate (Corning, 3513) at 400,000 cells per well in culture media. Cells were returned to 37 °C and 5% CO2 incubator to adhere to plates overnight. Days 1-4 the media contained DMEM, 10% FBS, 1% PenStrep with 10 μM Retinoic acid. Day 5-7 the media contained DMEM, 10% FBS, 1% PenStrep with 10μM Retinoic acid and 25 ng BDNF. Days 8-10 the media contained DMEM, 5% FBS, 1% PenStrep with 10 μM Retinoic acid and 25 ng BDNF. The final media for days 11-14 contained DMEM, 1% FBS, 1% PenStrep with 10 μM Retinoic acid and 25 ng BDNF.

**Western Blot**
For experiments without preformed fibrils, cells were plated into a 6-well plate (Corning, 3513) at 800,000 cells per well in culture media. Cells were returned to 37 °C and 5% CO2 incubator to adhere to plates overnight. Each well was treated with the indicated concentration of compound for the indicated time. Cells were washed with 1 mL PBS and harvested with trypsin. Cell pellets were washed with 1 mL PBS. Pellets were lysed with M-PER lysis buffer (Thermo, 78503) containing 1X HALT inhibitor cocktail (Thermo, 78442), and 125 U/mL benzonase (MilliporeSigma, 70664). Cells were lysed on ice for 1 hr. Lysates were clarified by centrifuging at 21,000 x g for 15 min at 4°C.

For experiments with preformed fibrils, HEK293 cells were plated into a 6-well plate (Corning, 3513) at 600,000 cells per well in culture media. Cells were returned to 37 °C and 5% CO2 incubator to adhere to plates overnight. SHSY5Y neurons were used for experiments on day 14 of differentiation. The following day, each well was transfected with 4 μg alpha-synuclein preformed fibrils, sonicated with probe sonicator directly before use, using either Lipofectamine 3000 or PEI star according to manufacturer instructions. Cells were returned to 37 °C and 5% $CO_2$ incubator for 16 hr. Media was discarded and cells were washed with 1 mL PBS three times to remove exogenous fibrils. Media was replaced with fresh media. Each well was treated with the indicated concentration of compound for the indicated time. Cells were washed twice with 1 mL PBS and harvested with trypsin. Cell pellets were washed with 1 mL PBS. Pellets were lysed with M-PER lysis buffer (Thermo, 78503) + 3% SDS containing 1X HALT inhibitor cocktail (Thermo, 78442), and 125 U/mL benzonase (MilliporeSigma, 70664). Cells were lysed on ice for 1 hr. Lysates were sonicated with Branson water bath sonicator for 10 min. Lysates were clarified by centrifuging at 21,000 x g for 15 min at 4°C.

For all experiments, protein concentration was determined by bicinchoninic acid (BCA) assay (Thermo, 23225). SDS-PAGE samples were prepared in 1X NuPage LDS sample buffer (Invitrogen, NP0007) and boiled at 95 °C for 5 min. SDS-PAGE samples were run on a Bolt 4–12% Bis-Tris Gel (Invitrogen, NW04125BOX) in MES run buffer (Invitrogen, B0002) for 50 min at 180 V, then transferred to a nitrocellulose membrane (Cytiva, 10600011) in Bolt transfer buffer (Invitrogen, BT00061) for 90 min at 45 V. Membranes were fixed with 0.4% (w/v) paraformaldehyde in water for 30 min at room temperature. Membrane was blocked with 5% non-fat dry milk (Kroger) in TBST (Thermo, 28360) for 1 hr at room temperature, then incubated with primary antibody diluted in TBS blocking buffer (LI-COR, 92760001) overnight at 4 °C. Primary antibodies used were 1:1,000 alpha-synuclein (Abcam, ab138501), 1:1,000 HA (Cell Signaling Technology, 3724S) and 1:1,000 beta-actin (Cell Signaling Technology, 3700S). Membrane was washed three times with TBST then incubated with DyLight 680 anti-mouse IgG (Cell Signaling Technology, 5470S) and DyLight 800 anti-rabbit IgG (Cell Signaling Technology, 5151S) diluted 1:10,000 in TBS blocking buffer for 1 hr at room temperature. Membrane was washed three times with TBST then imaged using a ChemiDoc Imaging System (Bio-Rad).

**Recombinant expression of alpha-synuclein constructs**

pET plasmid containing either wild type alpha-synuclein or minimal N degron and HA tagged alpha-synuclein was transformed into Rosetta (DE3) *E. coli* cells (Sigma, 70954). Seed cultures were grown from freshly transformed colonies and used to inoculate a 1% v/v ratio into Luria-Bertani medium with ampicillin (100 µg/mL). Cultures were subsequently grown at 37˚C while being shaken 200 rpm to an OD600 of 0.6-0.8 in which protein expression was induced with 0.5 mM isopropyl β-thiogalactopyranoside. Cell cultures were incubated for 4 hours at 37˚C and harvested by centrifugation at 5,000 xg at 4°C for 20 minutes. Cell pellets were either frozen at -80°C or immediately used for protein purification.

*Wild type alpha-synuclein:*
MDVFMKGLSKAKEGVVAAAEKTKQGVAEAAGKTKEGVLYVGSKTKEGVVHGVATVAEKTKEQ
VTNVGGAVVTGVTAVAQKTVEGAGSIAAATGFVKKDQLGKNEEGAPQEGILEDMPVDPDNEA
YEMPSEEGYQDYEPEA

*Minimal N degron tagged alpha-synuclein:*
MLQCEICGFTCRQKGNLLRHIKLHGGGGSGGGGSMDVFMKGLSKAKEGVVAAAEKTKQGVA
EAAGKTKEGVLYVGSKTKEGVVHGVATVAEKTKEQVTNVGGAVVTGVTAVAQKTVEGAGSIAA
ATGFVKKDQLGKNEEGAPQEGILEDMPVDPDNEAYEMPSEEGYQDYEPEAGYPYDVPDYA

**Purification of wild type and degron tagged alpha-synuclein**

Wild type and minimal N degron tagged alpha-synuclein were purified using the ammonium sulfate precipitation procedure described in the following reference with adaptations.[1] Briefly, the cell pellet was thawed and resuspended in lysis buffer (10 mM Tris, 1 mM EDTA pH 7.2) with Roche cOmpleteTM Protease Inhibitor Cocktail (Sigma, 11836170001) and lysed by probe sonication for 15 minutes using a Qsonica sonicator with a 1/8” diameter probe tip at 12 kHz (60%) output. The lysate was then clarified by centrifugation at 27,000 xg for 20 minutes and the supernatant was separated from the pellet and a saturated ammonium sulfate solution (4.1M) was added dropwise over a 20-minute period to a final ammonium sulfate saturation of 47% to precipitate the protein. The solution was allowed to stir for an additional 30 minutes and followed by centrifugation (27,000 xg for 30 minutes) to pellet the precipitated protein. The protein pellet was resuspended in 10 mM Tris and 1 mM ethylenediaminetetraacetic acid (EDTA) (pH 7.4) and dialyzed overnight. The next day the sample was filtered through a 0.22 µm filter and loaded onto a HiTrap Q HP anion exchange chromatography column (Cytiva). The protein was eluted using a linear gradient of 10 mM Tris (pH 7.5) to 10 column volumes of 10 mM Tris and 0.75 M NaCl (pH 7.5). Fractions containing alpha-synuclein were pooled and concentrated. 6 M guanidine hydrochloride, 5 mM tris(2-carboxyethyl)phosphine (TCEP), and 0.1% trifluoracetic acid (TFA) were added to the sample, and the sample was purified by reverse-phase high pressure liquid chromatography (HPLC) using an XBridge Peptide BEH C18 OBD prep column (130 Å pore size, 10 µm particle size, 19 mm X 250 mm) on a Waters pump system. HPLC purifications were done using a gradient method of water and acetonitrile with 0.1% TFA as the mobile phases. The sample identity was confirmed by intact mass TOFMS using an Agilent 1260 Infinity Binary LC coupled to a 6230 TOFMS system (Agilent Technologies, Santa Clara, CA). Samples were lyophilized until use.

**Protein aggregation**

Wild type and Minimal N degron alpha-synuclein fibrils were prepared from monomer protein at 6 mg/mL in aggregation buffer (10 mM sodium phosphate (pH 7.4), 150 mM NaCl, 0.5 mM TCEP and 0.01% $NaN_3$ buffer). The fibrilization reaction was carried out on a 200 µL scale and shaken at 400 rpm for 7 days. The fibrils were then centrifuged and washed twice with deionized water. The fibrils were then resuspended in 200 µL deionized water and bath sonicated for 30 minutes. The fibrils were then aliquoted in 20 µL aliquots and stored at -80 °C.

**ThT aggregation assay**

To characterize the aggregation kinetics of Minimal N degron alpha-synuclein, an aggregation kinetics assay was performed. Samples were prepared in 96-well plates with a total assay volume of 100 µL. Minimal N degron alpha-synuclein was prepared at a concentration of 50 µM in aggregation buffer with 20 µM Thioflavin T (THT) (Sigma, 596200). Preformed wild type alpha-synuclein fibrils were added at varying concentrations from 0 to 500 nM. The assay plate was incubated at 37 °C and THT emission was monitored at 480 nm (excitation wavelength of 440 nm) every 15 min with shaking for 30 s before reading. The assay was performed on a SpectraMax® M3 microplate reader (Molecular Devices, San Jose, CA). To determine aggregation kinetics parameters (kapp, t1/2, tlag) the THT curves were plotted and analyzed according to a sigmoidal equation as described in the following references.[2,3]

**Transmission electron microscopy (TEM)**

Wild type or degron tagged alpha-synuclein were loaded onto freshly glow-discharged Formvar/Carbon 400 mesh copper grids (Ted Pella, Inc.). Briefly, the protein was placed onto the grids in 10 µL drops and allowed to adsorb onto the surface for 5 minutes, the grids were then washed with water 3 times and stained with drops containing 1% (w/v) uranyl acetate for 1 min (Ladd Research Industries, Williston VT). The samples were left to dry for 20 min and stored overnight. Grid images were acquired using a JEOL JEM-1400Plus transmission electron microscope (80 kV) equipped with a Gatan OneView camera.

**Global proteomics profiling**

For experiments without preformed fibrils, cells were plated into a 6-well plate (Corning, 3513) at 800,000 cells per well in culture media. Cells were returned to 37 °C and 5% $CO_2$ incubator to adhere to plates overnight. Each well was treated with the indicated concentration of compound for the indicated time in triplicate. Cells were washed with 1 mL PBS and harvested with trypsin. Cell pellets were washed three times with 1 mL PBS. Pellets were lysed with M-PER lysis buffer (Thermo, 78503) containing 1X HALT inhibitor cocktail (Thermo, 78442), and 125 U/mL benzonase (MilliporeSigma, 70664). Cells were lysed on ice for 1 hr. Lysates were sonicated with a Branson water bath sonicator for 10 min. Lysates were clarified by centrifuging at 21,000 x g for 15 min at 4°C.

For experiments with preformed fibrils, cells were plated into a 6-well plate (Corning, 3513) at 600,000 cells per well in culture media. Cells were returned to 37 °C and 5% $CO_2$ incubator to adhere to plates overnight. The following day, each well was transfected with 4 µg alpha-synuclein preformed fibrils, sonicated with probe sonicator directly before use, using either Lipofectamine 3000 or PEI star according to manufacturer instructions. Cells were returned to 37 °C and 5% $CO_2$ incubator for 16 hr. Media was discarded and cells were washed with 1 mL PBS three times to remove exogenous fibrils. Media was replaced with fresh media. Each well was treated with the indicated concentration of compound for the indicated time in triplicate. Cells were washed twice with 1 mL PBS and harvested with trypsin. Cell pellets were washed two times with 1 mL PBS. Pellets were lysed with M-PER lysis buffer (Thermo, 78503) + 3% SDS containing 1X HALT inhibitor cocktail (Thermo, 78442), and 125 U/mL benzonase (MilliporeSigma, 70664). Cells were lysed on ice for 1 hr. Lysates were sonicated with a Branson water bath sonicator for 10 min. Lysates were clarified by centrifuging at 21,000 x g for 15 min at 4°C.

For all experiments, Pierce BCA kit (Thermo Scientific, 23225) was used to determine total protein concentration. For each sample, 200 µg of protein was aliquoted into a fresh lo-bind Eppendorf tube and adjusted to 1 µg/µL for proteomics preparation. Samples were reduced with 10 mM DTT (Invitrogen, A39255) at 56 °C with shaking for 30 minutes, alkylated using 40 mM iodoacetamide (Sigma-Aldrich, 1149-25G) in dark at room temperature for 15 minutes. Protein was precipitated via methanol chloroform method to remove detergent. Precooled methanol and chloroform was added to each sample at a 4:4:1 (vol/vol/vol) lysate/methanol/chloroform. Samples were centrifuged at 14,000 g for 5 min at 4°C. Liquid was aspirated and protein was incubated with precooled methanol and centrifuged at 14,000 g for 5 min at 4°C. The methanol was aspirated and one final incubation, spin and aspiration with methanol was performed. Precipitated proteins were dried with caps open. Samples were completely resuspended in 1 M Urea. The samples were then incubated with 2 µg trypsin (Promega, V5111) overnight at 37 °C. The samples were then acidified to pH <= 2 and desalted using Pierce desalting spin columns (Thermo Scientific, 89852). Eluted peptides were dried down using SpeedVac and resuspended in 0.1% formic acid.

LC-MS/MS data were acquired using an Orbitrap Eclipse Tribrid Mass Spectrometer (Thermo Scientific) coupled with a Vanquish Neo UHPLC system (Thermo Scientific). Each sample was dissolved in 0.1% formic acid in water and loaded onto a 75 µm inner diameter homemade microcapillary column packed with 20 cm of Bridged Ethylene Hybrid C18 particles (1.7 µm, 130 Å, Waters™), with an integrated emitter tip. Mobile phase A consisted of water with 0.1% formic acid, and mobile phase B consisted of 80% acetonitrile (ACN) with 0.1% formic acid. LC separation was performed over a 140-minute gradient elution of 3–40% mobile phase B at a flow rate of 300 nL/min in DIA mode. The resolution was set to 120,000 at m/z 200 across the m/z range of 350–1200. The MS1 normalized AGC target was set to 100%, with a maximum injection time set to auto. DIA MS2 scans were acquired over the 200–1500 m/z range with a normalized AGC target of 800% and a maximum injection time set to auto, using an HCD collision energy setting of 30% and a default charge state of +2.

Protein identification was carried out using DIA-NN 1.8.1[4], with an in silico spectral library generated from the human reference proteome (UP000005640_9606, version 2024-03-07). Default parameters were used, including a 1% precursor FDR, with the following exceptions: tryptic specificity (--cut K*, R*, !*P), up to one missed cleavages, and two variable modifications per peptide. Methionine oxidation and N-terminal acetylation were considered variable modifications. Mass accuracy was set to 10 ppm, and MS1 accuracy was also set to 10 ppm. The report.pg_matrix output file from DIA-NN was used for analysis. In brief, data was median normalized, log2 transformed and imputed using the missForest package[5] in R. The fold change and p-value (Student's T-test) were calculated in R. The mass spectrometry proteomics data have been deposited to the ProteomeXchange Consortium via the PRIDE[6] partner repository with the dataset identifier PXD074612.
